# A booster dose enhances immunogenicity of the COVID-19 vaccine candidate ChAdOx1 nCoV-19 in aged mice

**DOI:** 10.1101/2020.10.27.357426

**Authors:** Alyssa Silva-Cayetano, William S. Foster, Silvia Innocentin, Sandra Belij-Rammerstorfer, Alexandra J. Spencer, Oliver T. Burton, Sigrid Fra-Bidó, Jia Le Lee, Nazia Thakur, Carina Conceicao, Daniel Wright, Jordan Barett, Nicola Evans-Bailey, Carly Noble, Dalan Bailey, Adrian Liston, Sarah C. Gilbert, Teresa Lambe, Michelle A. Linterman

**Affiliations:** Lymphocyte Signalling and Development, Babraham Institute, Babraham Research Campus, Cambridge, CB22 3AT, United Kingdom; The Jenner Institute, University of Oxford, Old Road Campus Research Building, Roosevelt Drive, Oxford, OX3 7DQ, United Kingdom; Biological Services Unit, Babraham Institute, Babraham Research Campus, Cambridge, CB22 3AT, United Kingdom; The Pirbright Institute, Ash Road, Pirbright, GU24 0NF, United Kingdom

## Abstract

The spread of SARS-CoV-2 has caused a global pandemic that has affected almost every aspect of human life. The development of an effective COVID-19 vaccine could limit the morbidity and mortality caused by infection, and may enable the relaxation of social distancing measures. Age is one of the most significant risk factors for poor health outcomes after SARS-CoV-2 infection, therefore it is desirable that any new vaccine candidates should elicit a robust immune response in older adults. Here, we test the immunogenicity of the adenoviral vectored vaccine ChAdOx1 nCoV-19 (AZD-1222) in aged mice. We find that a single dose of this vaccine induces cellular and humoral immunity in aged mice, but at a reduced magnitude than in younger adult mice. Furthermore, we report that a second dose enhances the immune response to this vaccine in aged mice, indicating that a prime-boost strategy may be a rational approach to enhance immunogenicity in older persons.

## Introduction

The current COVID-19 pandemic is caused by the zoonotic severe acute respiratory syndrome coronavirus-2 (SARS-CoV-2)(*1, 2*). The pandemic has affected almost every aspect of human life, and will continue to do so until effective vaccines or therapeutics are developed. SARS-CoV-2 infection is initiated when the trimeric ‘spike’ glycoprotein on the virion surface binds angiotensin-converting enzyme 2, allowing viral entry and initating viral replication(*3*). After an asymptomatic incubation period, the infection can cause highly heterogenous clinical outcomes, from negligible or mild symptoms to critical disease resulting in death(*4*). One of the main risk factors for severe disease and death is *age*(*4–6*). Therefore, successful interventions to limit the severe health outcomes of COVID-19 disease should aim to be effective in older adults.

Vaccines have been one of the most successful medical interventions to date(*7, 8*) and represent a potential solution to control the COVID-19 pandemic(*9, 10*). However, age-related changes in the immune system mean that older individuals often do not generate protective immunity after vaccination(*11–14*). As older persons are more at risk of severe health outcomes after infection, significant effort has been made to specifically tailor vaccines to be more effective in people aged 65 years or older. Modifications to vaccines include increasing the antigen dose and the use of adjuvants, which can support then enhancement of immune responses in older people compared to standard formulations of vaccines(*15, 16*). This indicates that the success of a SARS-CoV-2 vaccine may require alterations to the dosing regimen, including an increase in the dose or the number of doses in order to induce a sufficient immune response in older people.

The effectiveness of COVID-19 vaccine candidates in older adults will ultimately be determined in clinical trials. However, studies in aged animals can be used to test alternative vaccine strategies or dosing regimens, and can therefore be used to inform clinical strategy. This pre-clinical work may therefore expedite the process of developing an effective vaccine against COVID-19 for older adults. Despite ageing occurring on different time scales in mice and people, many of the cellular and molecular changes that occur with ageing are conserved between the species(*17*), including those in the immune system such as thymic involution and the loss of naïve T cells(*18*). The response to vaccination is no exception(*19, 20*); after vaccination, both aged mice (>20 months old) and older humans have reduced vaccine-specific antibody formation, an impaired type I interferon response and fewer T follicular helper cells(*21–24*). This impaired immune response to vaccination in older mice and humans has been linked with reduced protection against subsequent infection(*25–28*). Importantly, interventions that enhance immunogenicity of vaccines in aged mice are also effective in humans(*21, 27, 29, 30*), demonstrating that aged mice are a relevant pre-clinical model for testing the immunogenicity of new vaccine candidates.

ChAdOx1 nCoV-19 is a chimpanzee adeno (ChAd)-vectored vaccine that encodes the full-length spike protein of SARS-CoV-2. Vaccination with ChAdOx1 nCoV-19 elicits robust cellular and humoral immunity in mice, pigs and macaques(*31, 32*). Importantly, a single vaccination prevents viral pneumonia and reduces the viral load in rhesus macaques after high-dose SARS-CoV-2 challenge(*31*). The phase I/II ChAdOx1 nCoV-19 trial demonstrated that this vaccine has an acceptable safety profile and is immunogenic in adults between 18 and 55 years of age, stimulating virus-neutralising antibodies and spike-reactive T cells(*33*). In macaques, pigs and people, a homologous booster dose of ChAdOx1 nCoV-19 increased anti-spike antibody responses, indicating this is a safe and effective strategy to enhance immunogenicity(*31–33*).

Here, we demonstrate that a single dose of ChAdOx1 nCoV-19 elicits a B and T cell response in 3-month-old adult mice, with formation of plasma cells, germinal centres and T follicular helper cells contributing to anti-spike antibody production. The development of humoral immunity is complemented by the formation of polyfunctional vaccine-specific Th1 cells and CD8^+^ T cells. In aged 22-month-old mice a single dose of ChAdOx1 nCoV-19 induced the formation of Th1 cells, vaccine-reactive CD8^+^ T cells, a germinal centre response and vaccine-specific antibodies. However, the cellular and humoral response was reduced in magnitude in 22-month-old mice compared to 3-month-old adult mice, with antibody isotypes and subclasses produced being of a similar profile. Administration of a second dose enhanced the germinal centre response and antibody titre in aged mice, and also boosted the numbers of granzyme B producing CD8^+^ T cells. Together, this indicates that the immunogenicity of ChAdOx1 nCoV-19 can be enhanced in older individuals through the use of a prime-boost vaccination strategy.

## Results

### Intramuscular immunisation with ChAdOx1 nCoV-19 activates antigen presenting cells in the aortic lymph node and spleen

To map where antigen drains upon intramuscular immunisation, we immunised mice with fluorescent nanoparticles that freely drain into lymphoid organs(*34*). The nanoparticles were injected into the right quadriceps muscle of 3-month-old adult mice. Twenty-four hours after injection, confocal microscopy revealed fluorescent nanoparticles in the aortic lymph node (aLN) and the spleen of all immunised mice (**Fig. 1a and Sup. Fig. 1a**), while there was negligible detection of fluorescent nanoparticles in the inguinal and popliteal lymph nodes (data not shown). These data suggest the aLN and spleen are the main sites of antigen drainage upon intramuscular immunisation. As such, this study focused on the immune response induced by the ChAdOx1 nCoV-19 vaccine in these two immune compartments.

**Figure 1.**
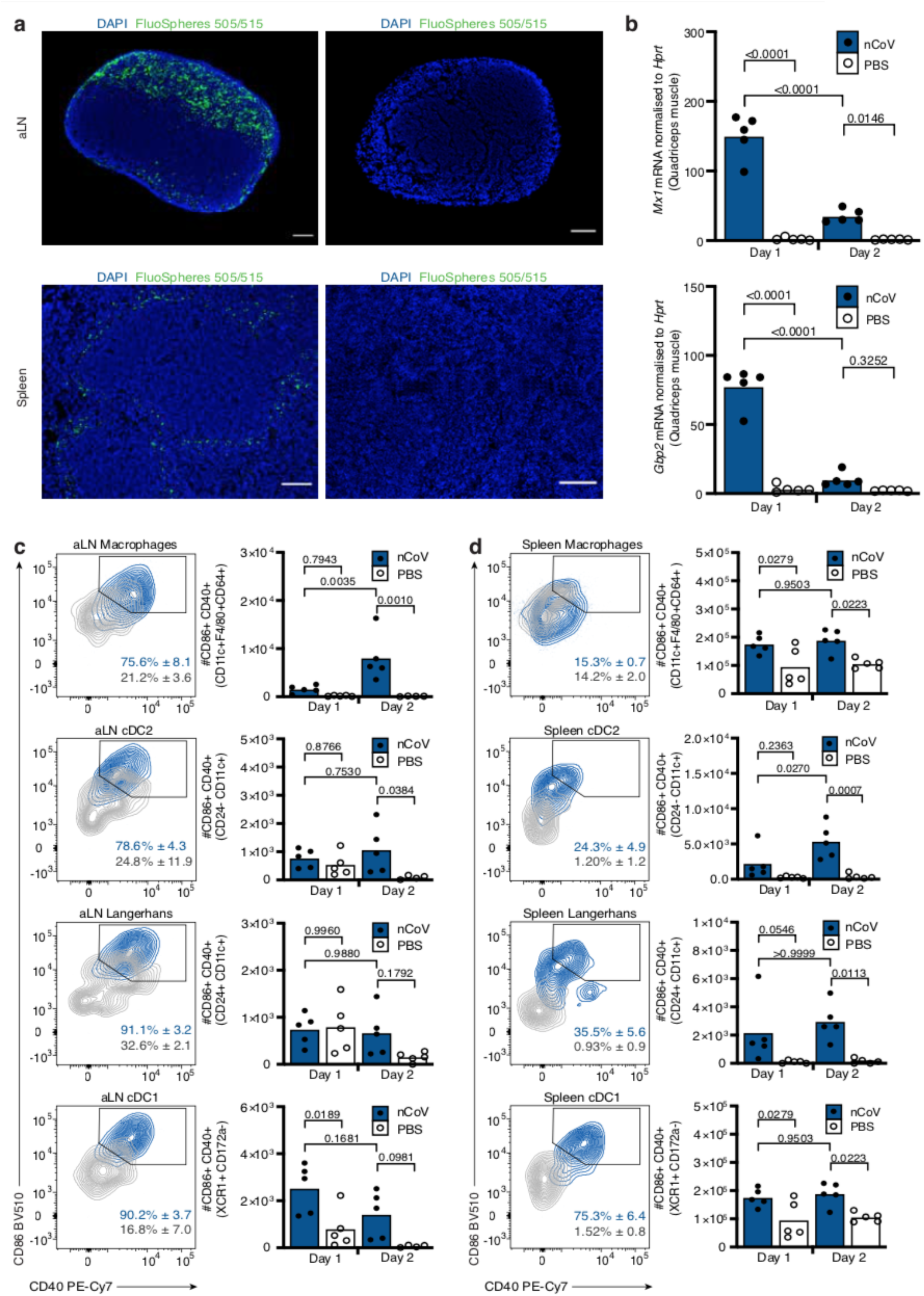
Intramuscular immunisation results in the activation of antigen presenting cells in the aLN and spleen upon immunisation with ChAdOx1 nCoV-19. **a.** Representative immunofluorescence confocal images of DAPI expression and FluoSpheres™ (505/515) localisation in the aLN (top) and spleen (bottom) of mice immunised with yellow-green 20nm fluorescent FluoSpheres™ (left) or PBS (right) at 24hr post intramuscular immunisation. Scale bars, 100μM for both aLN and spleen. Data are representative of two independent experiments (n=2-5 per group/experiment). **b.** Expression of *Mx1* (top) and *Gbp2* (bottom) in the quadriceps muscle of mice immunised with ChAdOx1 nCoV or PBS at one- and two-days post immunisation, as determined by RT-qPCR. **c-d.** Quantification of activated CD86^+^CD40^+^ macrophages (CD11c^+^F4/80^+^CD64^+^), cDC2 (CD24^-^CD11c^+^CD172a^+^XCR1^-^), Langerhan cells (CD24^+^CD11c^+^CD172a^+^XCR1^-^) and cDC1 (XCR1^+^CD172a^-^) in the aLN (**c**) and spleen (**d**) of mice immunised with ChAdOx1 nCoV or PBS at one-and two-days post i.m. immunisation. Representative flow cytometry plots (left) indicate the percentage, median and standard deviation of CD86^+^CD40^+^ cells for each antigen presenting cell population in mice immunised with ChAdOx1 nCoV-19 (blue) or PBS (grey). Quantification of the absolute number of CD86^+^CD40^+^ cells for each APC population is shown on bar graphs (right). Bar height in **b-d** corresponds to the mean and each circle represents one biological replicate. P-values were determined using a Shapiro-Wilk normality test followed by either an ordinary one-way ANOVA test for data with a normal distribution or a Kruskal Wallis test for non-normally distributed data alongside a multiple comparisons test. Data are representative of two independent experiments (n=4-5 per group/experiment).

To characterise the early events of the immune response to ChAdOx1 nCoV-19 vaccination, we immunised adult C57BL/6 mice and assessed the induction of interferon signalling as well as the expansion of antigen presenting cell populations at day 1 and 2 post immunisation. At the site of immunisation, the right quadriceps muscle, expression of the type I interferon inducible gene *Mx1* and the type 2 interferon inducible gene *Gbp2* was detected at day 1 in ChAdOx1 nCoV immunised mice compared to phosphate buffered saline (PBS) immunised controls (**Fig. 1b**). The expression of both genes was significantly reduced by day 2, indicating that the ChAdOx1 nCoV-19 vaccine induces a short-lived interferon response at the site of injection. In the draining aLN and spleen, the expansion and activation of macrophages, type 2 conventional dendritic cells (cDC2s), Langerhans cells and cDC1s was quantified by flow cytometry (**Sup. Fig. 1b**; gating strategy from (*35*)). In both the aLN and spleen of ChAdOx1 nCoV immunised mice there was a significant increase in the total number of CD11 c^+^ macrophages at day 2, while the total number of cDC2s, Langerhans and cDC1 s remained unperturbed compared to PBS-immunised controls (**Sup. Fig. 1c, d**). However, two days after immunisation, there was an increase in the total number of activated CD40^+^CD86^+^ cells in all antigen presenting cell populations compared to PBS-immunised controls, in both the aLN and spleen (**Fig. 1c, d**). The induction of adaptive immunity through T cell priming relies on the activation of these professional APCs; of which cDC2s have been implicated as the main subset to activate Tfh cells while the cDC1 subset is able to cross-present exogenous antigens to activate CD8^+^ T cells(*21, 36, 37*). These data demonstrate that the ChAdOx1 nCoV-19 vaccine activates several antigen presenting cell populations required for the initiation of cell-mediated and humoral immunity.

### ChAdOx1 nCoV-19 immunisation stimulates B cell activation and differentiation

To assess the adaptive immune response to ChAdOx1 nCoV-19 in detail we used a high-dimensional flow cytometry panel that contained antibodies that recognise multiple molecules used to define different lymphocyte subsets and their activation status (Table 1). Mice were immunised with either ChAdOx1 nCoV-19 or PBS, and the immune response was assessed seven, fourteen and twenty-one days later. t-distributed Stochastic Neighbour Embedding (tSNE) and FlowSOM analysis grouped B cells into five clusters (**Fig. 2a**). A cluster that corresponded to naïve B cells which were present in both ChAdOx1 nCoV-19 and PBS immunised mice, and four additional clusters which were overrepresented in ChAdOx1 nCoV-19 immunised mice. These clusters are CD69^+^ and CD86^+^ activated B cells, CD138^+^IRF4^+^ antibody-secreting plasma cells and Bcl6^+^GL-7^+^ germinal centre B cells. Manual biaxial gating of these populations (**Sup. Fig. 2a**) showed that ChAdOx1 nCoV-19 immunisation induced a robust B cell response in the draining lymph node and spleen (**Fig. 2b, c**), with early B cell activation and plasma cell formation occurring in the first week after immunisation, and the germinal centre response persisting over the three weeks assessed (**Fig. 2b-g**). Formation of T follicular helper and T follicular regulatory cells accompanied the germinal centre response (**Fig. 2h, i**). Antibodies binding the SARS-CoV-2 spike protein were induced by vaccination, and, as expected, the temporal induction of anti-spike IgM was faster than that of IgG, and serum IgA antibodies were not observed at high titre (**Fig. 2j**). A mix of anti-spike IgG antibody subtypes was observed, with IgG_2_ and IgG_1_ subclasses persisting at later timepoints. At all timepoints a predominantly Th1 dominant response (IgG_2_) was measured (**Fig. 2k**). The mean (and standard deviation) value for ratio of IgG_2_/IgG_1_ on day seven was 4.8 (1.4), on day fourteen 2.3 (0.5), and on day twenty-one 2.8 (1.8). This demonstrates that ChAdOx1 nCoV-19 immunisation stimulates B cell activation and differentiation, culminating in the production of anti-spike antibodies.

**Figure 2.**
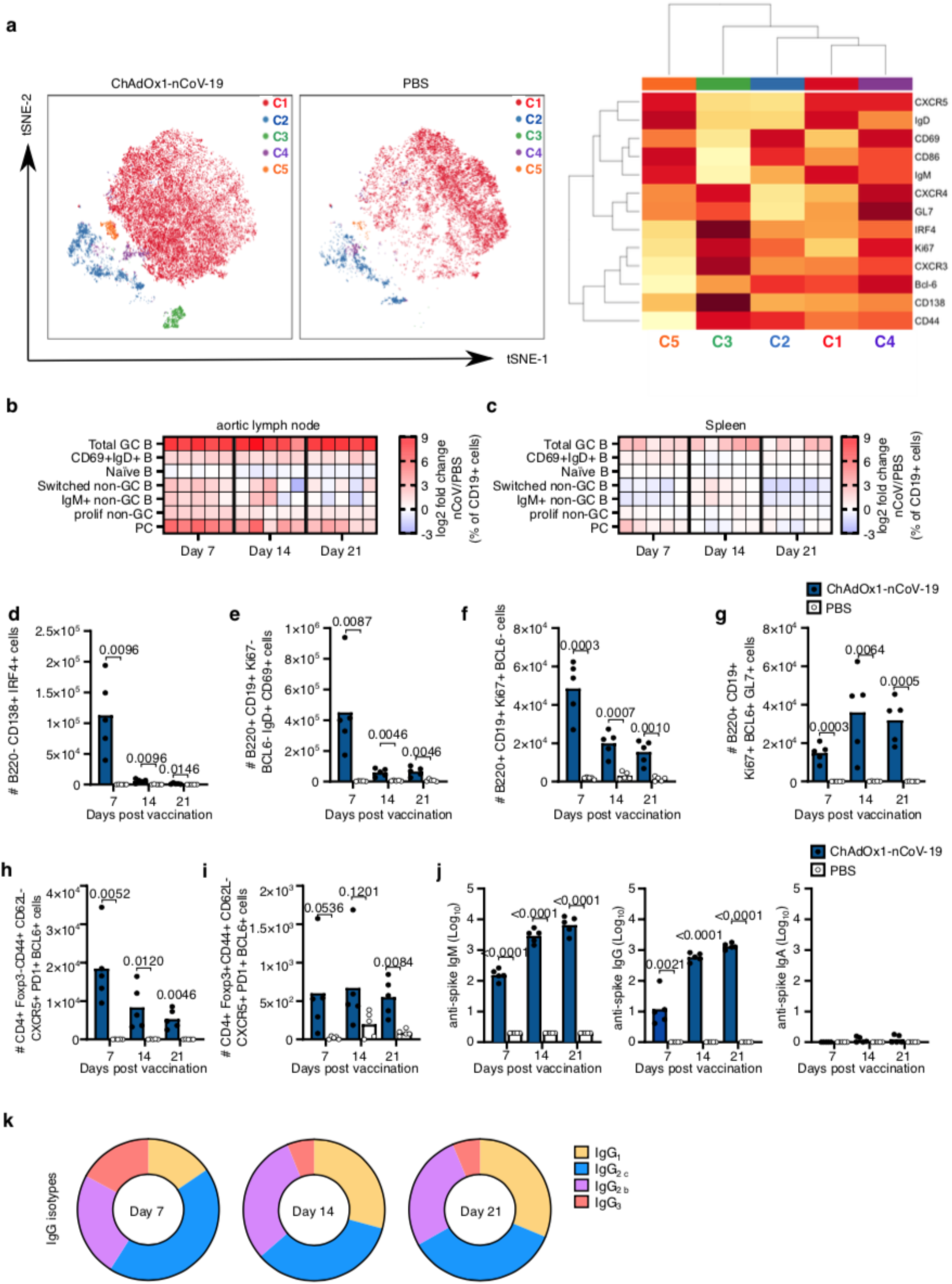
ChAdOx1 nCoV-19 induces a plasma cell and germinal centre B cell response. **a.** tSNE/FlowSOM analyses of CD19^+^ B cells from 3-month-old (3mo) mice seven days after immunization with ChAdOx1 nCoV-19 or PBS. Heatmaps of the manually gated B cell populations indicated at seven, 14 and 21 days after immunisation in the aortic lymph node (**b**) and spleen (**c**). Here the frequency of each cell subset in each ChAdOx1 nCoV-19 immunised mouse has been expressed as the log2 fold change over the average frequency in PBS immunised mice (n=5). Bar charts showing the total number of plasma cells (**d**), CD69^+^IgD^+^ B cells (**e**), proliferating non-germinal centre B cells (**f**) and germinal centre B cells (**g**) at the indicated time points after immunization. Number of T follicular helper (**h**) and T follicular regulatory cells (**i**) at the indicated time points post-immunisation. Serum anti-spike IgM, IgG, and IgA (**j**) antibodies seven, 14 and 21 days after immunization. **k**. Pie charts indicating the mean abundance of each IgG antibody subclass in the serum at the indicated time points after immunisation. In **d-j** bar height in corresponds to the mean and each circle represents one biological replicate. P-values are calculated using a student’s t-test with Holm-Sidak multiple testing correction, for ELISA data analyses were done on log transformed values.

**Table 1.**
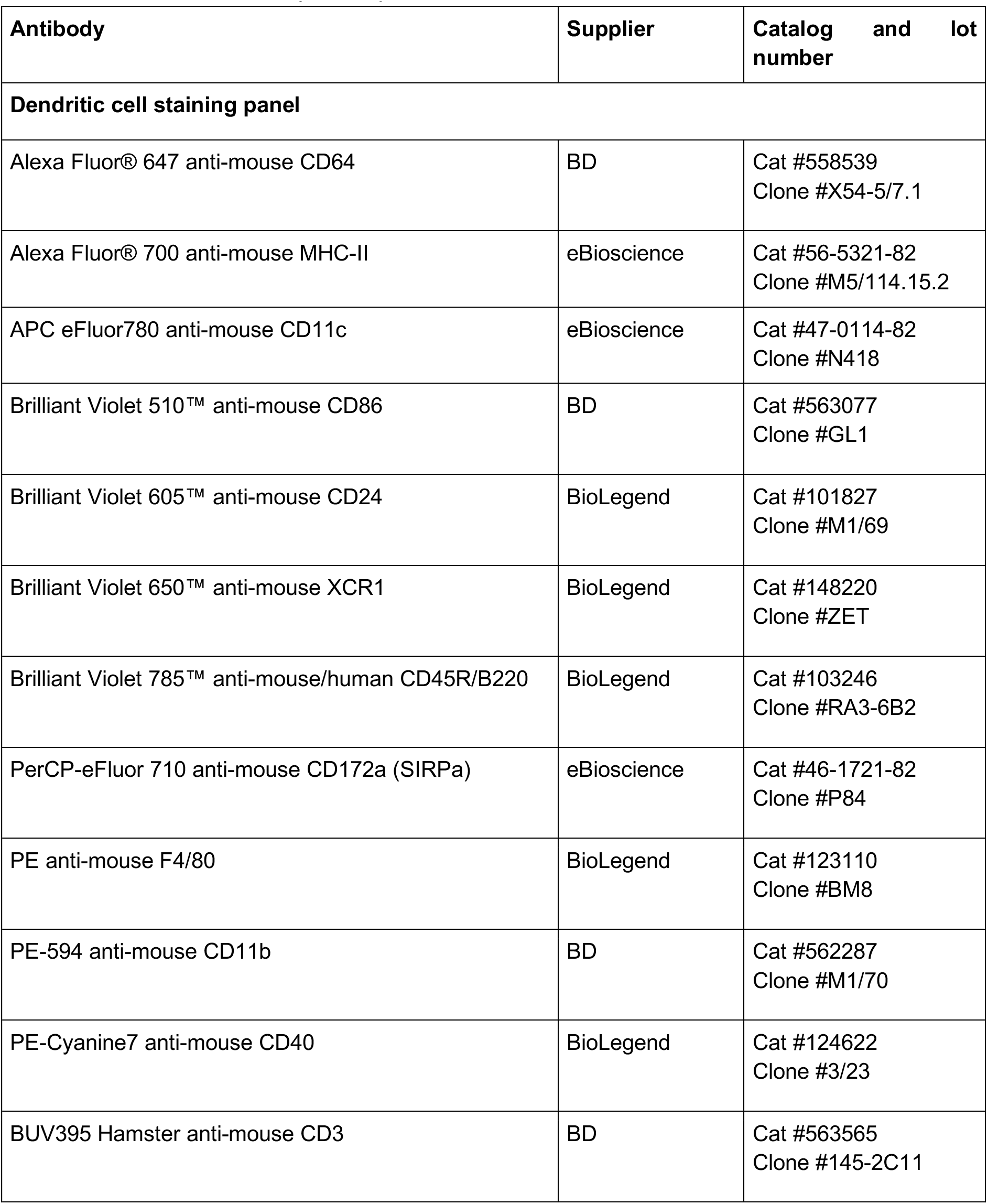
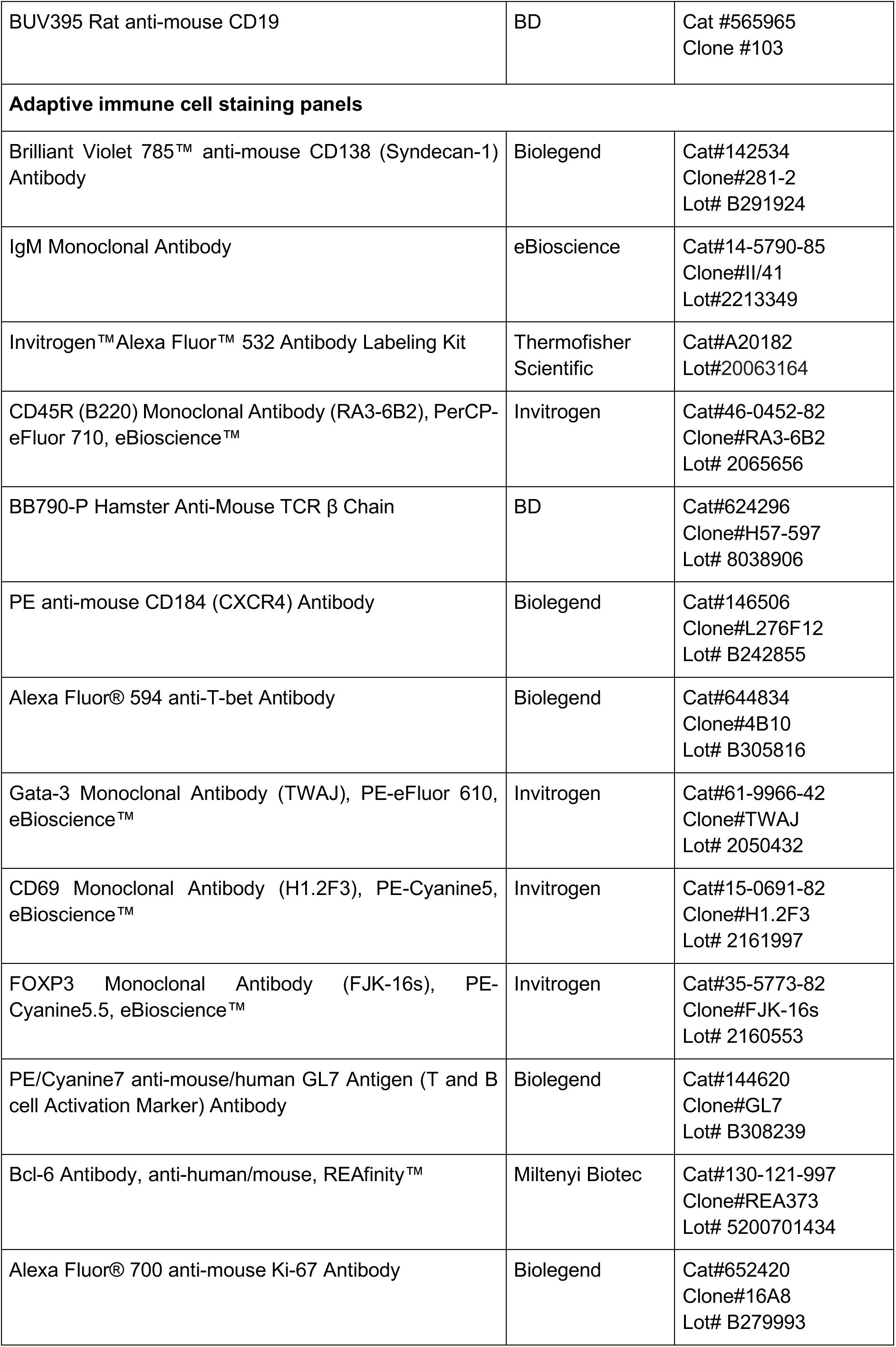
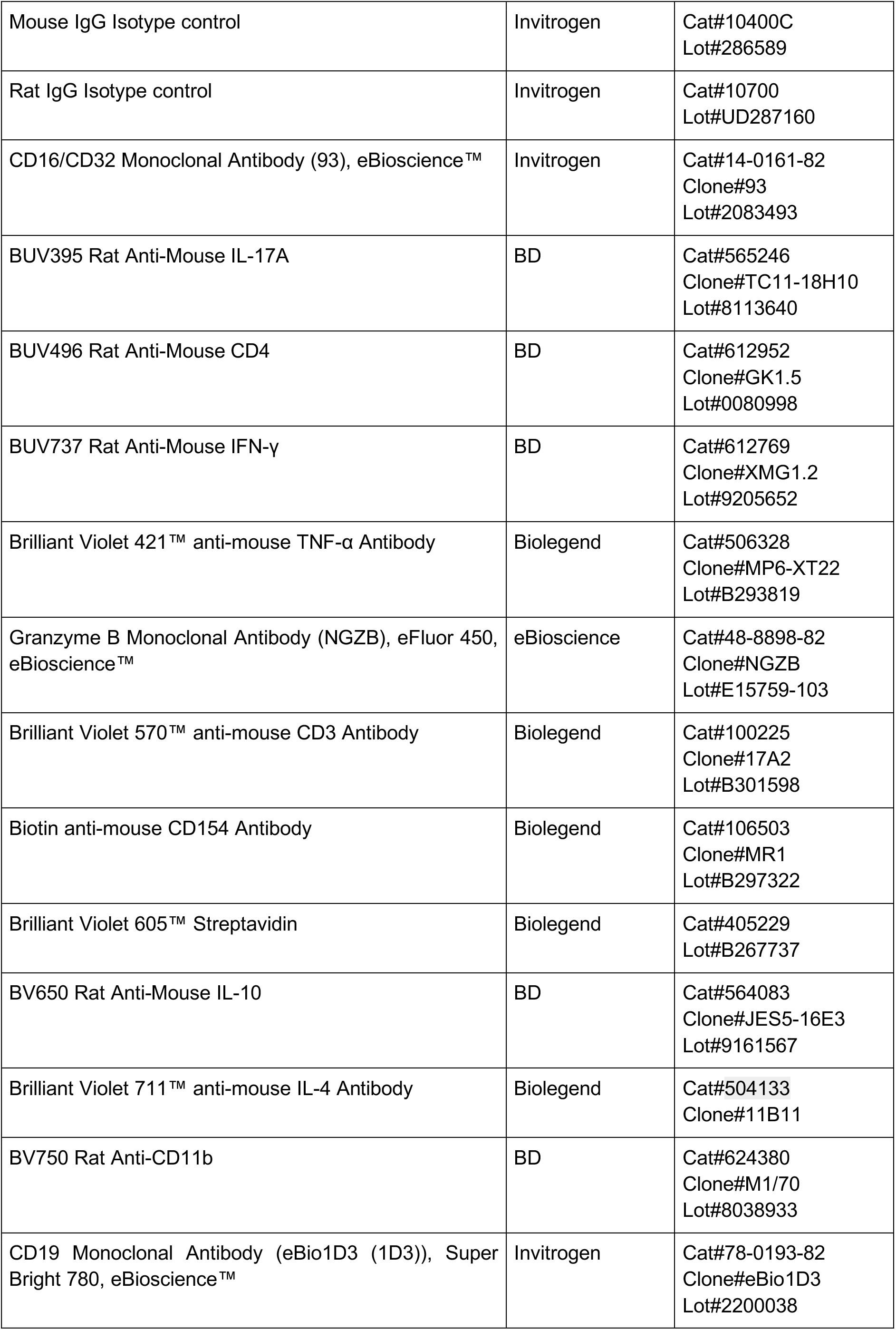
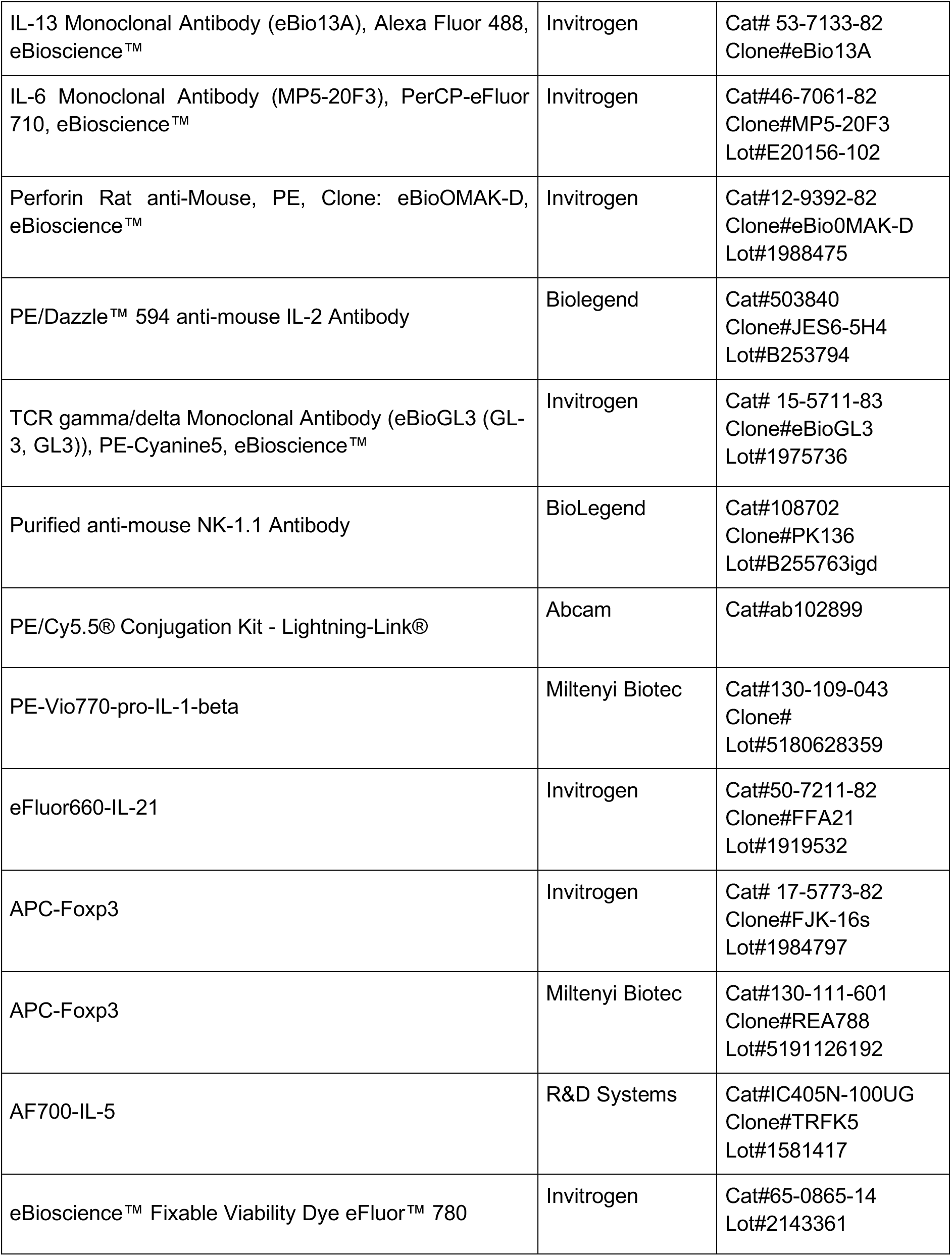
Antibodies for flow cytometry

| Antibody | Supplier | Catalog and lot number |
| --- | --- | --- |
| **Dendritic cell staining panel** |  |  |
| Alexa Fluor® 647 anti-mouse CD64 | BD | Cat #558539<br>Clone #X54-5/7.1 |
| Alexa Fluor® 700 anti-mouse MHC-II | eBioscience | Cat #56-5321-82<br>Clone #M5/114.15.2 |
| APC eFluor780 anti-mouse CD11c | eBioscience | Cat #47-0114-82<br>Clone #N418 |
| Brilliant Violet 510™ anti-mouse CD86 | BD | Cat #563077<br>Clone #GL1 |
| Brilliant Violet 605™ anti-mouse CD24 | BioLegend | Cat #101827<br>Clone #M1/69 |
| Brilliant Violet 650™ anti-mouse XCR1 | BioLegend | Cat #148220<br>Clone #ZET |
| Brilliant Violet 785™ anti-mouse/human CD45R/B220 | BioLegend | Cat #103246<br>Clone #RA3-6B2 |
| PerCP-eFluor 710 anti-mouse CD172a (SIRPa) | eBioscience | Cat #46-1721-82<br>Clone #P84 |
| PE anti-mouse F4/80 | BioLegend | Cat #123110<br>Clone #BM8 |
| PE-594 anti-mouse CD11b | BD | Cat #562287<br>Clone #M1/70 |
| PE-Cyanine7 anti-mouse CD40 | BioLegend | Cat #124622<br>Clone #3/23 |
| BUV395 Hamster anti-mouse CD3 | BD | Cat #563565<br>Clone #145-2C11 |
| BUV395 Rat anti-mouse CD19 | BD | Cat #565965<br>Clone #103 |
| **Adaptive immune cell staining panels** |  |  |
| Brilliant Violet 785™ anti-mouse CD138 (Syndecan-1) Antibody | Biolegend | Cat#142534<br>Clone#281-2<br>Lot# B291924 |
| IgM Monoclonal Antibody | eBioscience | Cat#14-5790-85<br>Clone#II/41<br>Lot#2213349 |
| Invitrogen™ Alexa Fluor™ 532 Antibody Labeling Kit | Thermofisher Scientific | Cat#A20182<br>Lot#20063164 |
| CD45R (B220) Monoclonal Antibody (RA3-6B2), PerCP-eFluor 710, eBioscience™ | Invitrogen | Cat#46-0452-82<br>Clone#RA3-6B2<br>Lot# 2065656 |
| BB790-P Hamster Anti-Mouse TCR β Chain | BD | Cat#624296<br>Clone#H57-597<br>Lot# 8038906 |
| PE anti-mouse CD184 (CXCR4) Antibody | Biolegend | Cat#146506<br>Clone#L276F12<br>Lot# B242855 |
| Alexa Fluor® 594 anti-T-bet Antibody | Biolegend | Cat#644834<br>Clone#4B10<br>Lot# B305816 |
| Gata-3 Monoclonal Antibody (TWAJ), PE-eFluor 610, eBioscience™ | Invitrogen | Cat#61-9966-42<br>Clone#TWAJ<br>Lot# 2050432 |
| CD69 Monoclonal Antibody (H1.2F3), PE-Cyanine5, eBioscience™ | Invitrogen | Cat#15-0691-82<br>Clone#H1.2F3<br>Lot# 2161997 |
| FOXP3 Monoclonal Antibody (FJK-16s), PE-Cyanine5.5, eBioscience™ | Invitrogen | Cat#35-5773-82<br>Clone#FJK-16s<br>Lot# 2160553 |
| PE/Cyanine7 anti-mouse/human GL7 Antigen (T and B cell Activation Marker) Antibody | Biolegend | Cat#144620<br>Clone#GL7<br>Lot# B308239 |
| Bcl-6 Antibody, anti-human/mouse, REAfinity™ | Miltenyi Biotec | Cat#130-121-997<br>Clone#REA373<br>Lot# 5200701434 |
| Alexa Fluor® 700 anti-mouse Ki-67 Antibody | Biolegend | Cat#652420<br>Clone#16A8<br>Lot# B279993 |
| Mouse IgG Isotype control | Invitrogen | Cat#10400C<br>Lot#286589 |
| Rat IgG Isotype control | Invitrogen | Cat#10700<br>Lot#UD287160 |
| CD16/CD32 Monoclonal Antibody (93), eBioscience™ | Invitrogen | Cat#14-0161-82<br>Clone#93<br>Lot#2083493 |
| BUV395 Rat Anti-Mouse IL-17A | BD | Cat#565246<br>Clone#TC11-18H10<br>Lot#8113640 |
| BUV496 Rat Anti-Mouse CD4 | BD | Cat#612952<br>Clone#GK1.5<br>Lot#0080998 |
| BUV737 Rat Anti-Mouse IFN-γ | BD | Cat#612769<br>Clone#XMG1.2<br>Lot#9205652 |
| Brilliant Violet 421™ anti-mouse TNF-α Antibody | Biolegend | Cat#506328<br>Clone#MP6-XT22<br>Lot#B293819 |
| Granzyme B Monoclonal Antibody (NGZB), eFluor 450, eBioscience™ | eBioscience | Cat#48-8898-82<br>Clone#NGZB<br>Lot#E15759-103 |
| Brilliant Violet 570™ anti-mouse CD3 Antibody | Biolegend | Cat#100225<br>Clone#17A2<br>Lot#B301598 |
| Biotin anti-mouse CD154 Antibody | Biolegend | Cat#106503<br>Clone#MR1<br>Lot#B297322 |
| Brilliant Violet 605™ Streptavidin | Biolegend | Cat#405229<br>Lot#B267737 |
| BV650 Rat Anti-Mouse IL-10 | BD | Cat#564083<br>Clone#JES5-16E3<br>Lot#9161567 |
| Brilliant Violet 711™ anti-mouse IL-4 Antibody | Biolegend | Cat#504133<br>Clone#11B11 |
| BV750 Rat Anti-CD11b | BD | Cat#624380<br>Clone#M1/70<br>Lot#8038933 |
| CD19 Monoclonal Antibody (eBio1D3 (1D3)), Super Bright 780, eBioscience™ | Invitrogen | Cat#78-0193-82<br>Clone#eBio1D3<br>Lot#2200038 |
| IL-13 Monoclonal Antibody (eBio13A), Alexa Fluor 488, eBioscience™ | Invitrogen | Cat# 53-7133-82<br>Clone#eBio13A |
| IL-6 Monoclonal Antibody (MP5-20F3), PerCP-eFluor 710, eBioscience™ | Invitrogen | Cat#46-7061-82<br>Clone#MP5-20F3<br>Lot#E20156-102 |
| Perforin Rat anti-Mouse, PE, Clone: eBioOMAK-D, eBioscience™ | Invitrogen | Cat#12-9392-82<br>Clone#eBioOMAK-D<br>Lot#1988475 |
| PE/Dazzle™ 594 anti-mouse IL-2 Antibody | Biolegend | Cat#503840<br>Clone#JES6-5H4<br>Lot#B253794 |
| TCR gamma/delta Monoclonal Antibody (eBioGL3 (GL-3, GL3)), PE-Cyanine5, eBioscience™ | Invitrogen | Cat# 15-5711-83<br>Clone#eBioGL3<br>Lot#1975736 |
| Purified anti-mouse NK-1.1 Antibody | BioLegend | Cat#108702<br>Clone#PK136<br>Lot#B255763igd |
| PE/Cy5.5® Conjugation Kit - Lightning-Link® | Abcam | Cat#ab102899 |
| PE-Vio770-pro-IL-1-beta | Miltenyi Biotec | Cat#130-109-043<br>Clone#<br>Lot#5180628359 |
| eFluor660-IL-21 | Invitrogen | Cat#50-7211-82<br>Clone#FFA21<br>Lot#1919532 |
| APC-Foxp3 | Invitrogen | Cat# 17-5773-82<br>Clone#FJK-16s<br>Lot#1984797 |
| APC-Foxp3 | Miltenyi Biotec | Cat#130-111-601<br>Clone#REA788<br>Lot#5191126192 |
| AF700-IL-5 | R&D Systems | Cat#IC405N-100UG<br>Clone#TRFK5<br>Lot#1581417 |
| eBioscience™ Fixable Viability Dye eFluor™ 780 | Invitrogen | Cat#65-0865-14<br>Lot#2143361 |

### ChAdOx1 nCoV-19 immunisation induces Th1 cell and Th1-like Treg formation

To investigate how ChAdOx1 nCoV-19 immunisation affects CD4^+^ T cells beyond the germinal centre-associated subsets, tSNE analysis followed by manual gating was performed (**Fig. 3a-c, Sup. Fig. 2b, c**). tSNE/FlowSOM clustering showed that vaccination induced populations of T cells that express markers associated with T follicular helper cells and Th1 cells (**Fig. 3a**). The conventional CD4^+^ T cell response to ChAdOx1 nCoV-19 was characterised by early activation and proliferation, as well as the formation of CXCR3^+^ Th1 cells (**Fig. 3a-f**). The Foxp3^+^ regulatory T cell response was characterised by early expression of Ki67, indicative of proliferation, and the differentiation of CXCR3^+^ Th1-like cells (**Fig. 3c, g**). To further characterise CD4^+^ T cell responses after vaccination, cells from the aLN were restimulated with PdBu/ionomycin and cytokine production was assessed. IL-2, TNFα, IL-10 and IFNγ were induced by cells isolated from animals that had been ChAdOx1 nCoV-19 vaccinated, with no difference observed in Th17-associated IL-17 or the Th2-associated cytokines IL-4 and IL-5 (**Fig. 3h, Sup. Fig. 3a, b**). Furthermore, restimulation of splenocytes with peptide pools from SARS-CoV-2 identified antigen-specific IL-2, TNFα, and IFNγ producing cells, with triple producers persisting 21-days after immunisation (**Fig. 3i, Sup. Fig. 3c**), consistent with previous work(*37*). Therefore, ChAdOx1 nCoV-19 vaccination induces an early formation of Th1 cells and Th1-like Treg cells accompanied by the induction of virus-specific Th1-skewed cytokine-secreting cells.

**Figure 3.**
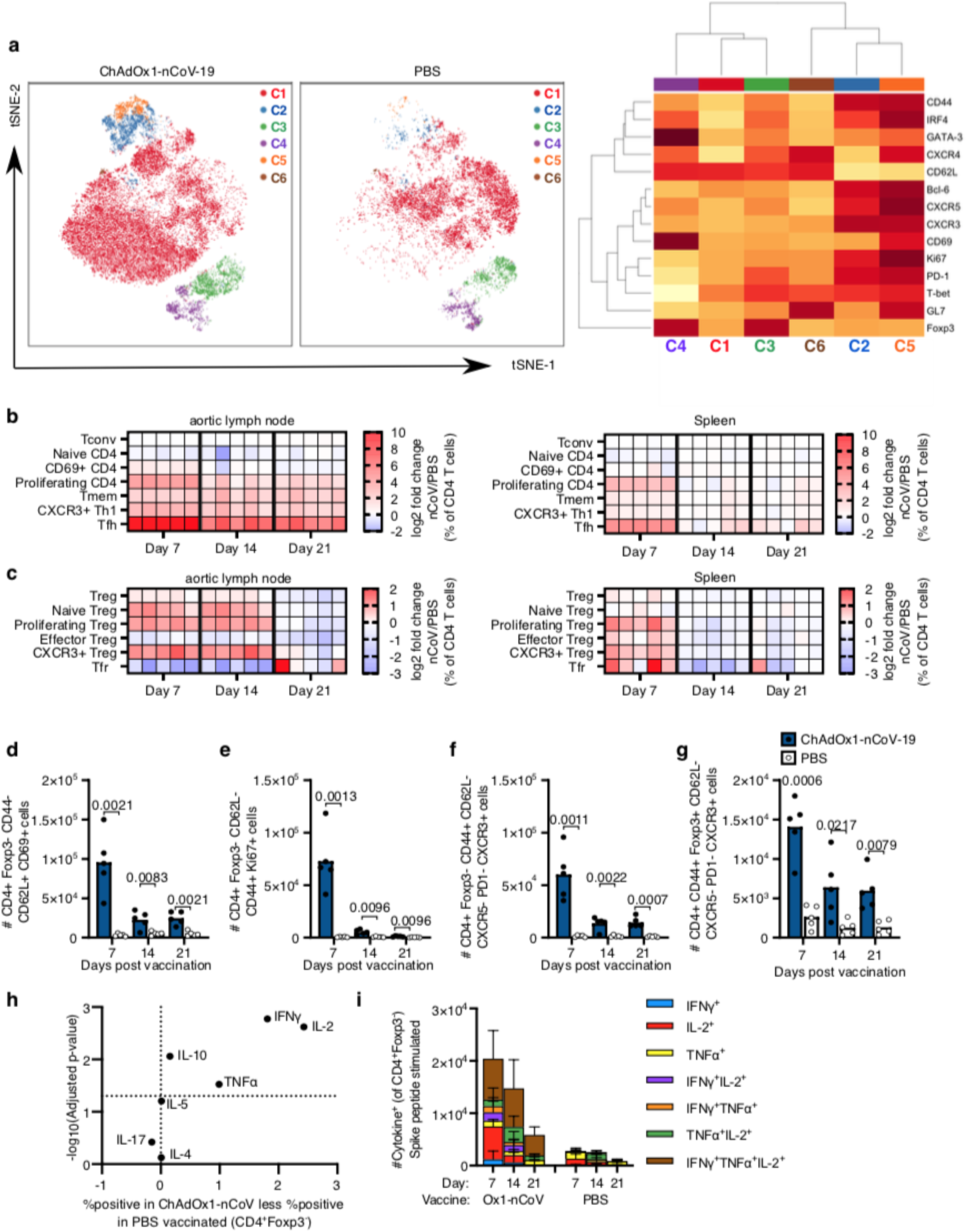
ChAdOx1 nCoV-19 induces a Th1 dominated CD4 cell response. **a.** tSNE/FlowSOM analyses of CD4^+^ T cells from 3-month-old (3mo) mice seven days after immunization with ChAdOx1 nCoV-19 or PBS. Heatmaps of the manually gated CD4^+^Foxp3^-^ (**b**) and Foxp3^+^CD4^+^ (**c**) T cell populations indicated at seven, 14 and 21 days after immunisation in the aortic lymph node (right) and spleen (left). Here the frequency of each cell subset in each ChAdOx1 nCoV-19 immunised mouse has been expressed as the log2 fold change over the average frequency in PBS immunised mice (n=5). Bar charts showing the number of CD69^+^CD62L^+^CD44^-^CD4^+^Foxp3^-^ (**d**), Ki67^+^CD4^+^Foxp3^-^ (**e**) CXCR3^+^ non-Tfh cells (**f**) and CXCR3^+^ Th1-like Treg cells (**g**) CD4^+^ cells in the aortic lymph node of ChAdOx1 nCoV-19 or PBS immunised mice, at the indicated timepoints post immunisation. **h**. Analysis of cytokine production six hours after PdBu/ionomycin stimulation of aortic lymph node cells from 3-month-old mice seven days after immunization with ChAdOx1 nCoV-19 or PBS. **i**. Stacked bar plots show the number of CD4^+^Foxp3^-^ cells singly or co-producing IFNγ, IL-2 or TNFα six hours after restimulation with SARS-CoV-2 peptide pools, each bar segment represents the mean and the error bars the standard deviation. In **d-g** bar height in corresponds to the mean and each circle represents one biological replicate. P-values are calculated using a student’s t-test with Holm-Sidak multiple testing correction.

### ChAdOx1 nCoV-19 induces a CD8^+^ T cell response

tSNE analysis of CD8^+^ T cells from the aLNs of ChAdOx1 nCoV-19 immunised and PBS control mice revealed distinct clustering seven days after immunisation. CD8^+^ T cell clusters that were present in ChAdOx1 nCoV-19 immunised mice, but not in PBS immunised animals expressed high levels of CD44, Ki67, CXCR3, PD-1, IRF4 or CD69, indicative of CD8^+^ T cell activation (**Fig. 4a**). In order to understand how the response evolves over time, manual gating of different CD8^+^ T cell populations (**Sup. Fig. 2d**), including those identified in the tSNE analysis, was done on samples taken seven, fourteen and twenty-one days after immunisation. In the aLN and spleen, the CD8^+^ T cell response to ChAdOx1 nCoV-19 was characterised by an increase in Ki67 expression, the upregulation of the activation markers CD69, CXCR3 and PD-1, as well as the formation of CD44^+^CD62L^-^ T effector memory cells (**Fig. 4b-f**). To characterise the overall production of granzyme B and cytokines by CD8^+^ T cells seven days after vaccination, cells from the aLN were restimulated with PdBu/ionomycin. There was a significant production of granzyme B, IL-2, TNFα and IFNγ in ChAdOx1 nCOV-19 vaccinated mice compared to PBS immunised mice (**Fig. 4g**). Restimulation of splenocytes with spike protein peptide pools from SARS-CoV-2 showed that antigen-specific cytokine producing CD8^+^ T cells form in response to immunisation (**Fig. 4g, Sup. Fig. 4a, b**). Granzyme B producing CD8^+^ T cells formed early, and tended not to co-produce cytokines (**Fig. 4h**). Both single and multiple cytokine producing cells formed at all time points, with TNFα and IFNγ being the dominant cytokines expressed (**Fig. 4h, Sup. Fig. 4c**). With this dosing regimen, this data demonstrates that ChAdOx1 nCoV-19 stimulates a robust CD8^+^ T cell response that peaks around the first week after vaccination.

**Figure 4.**
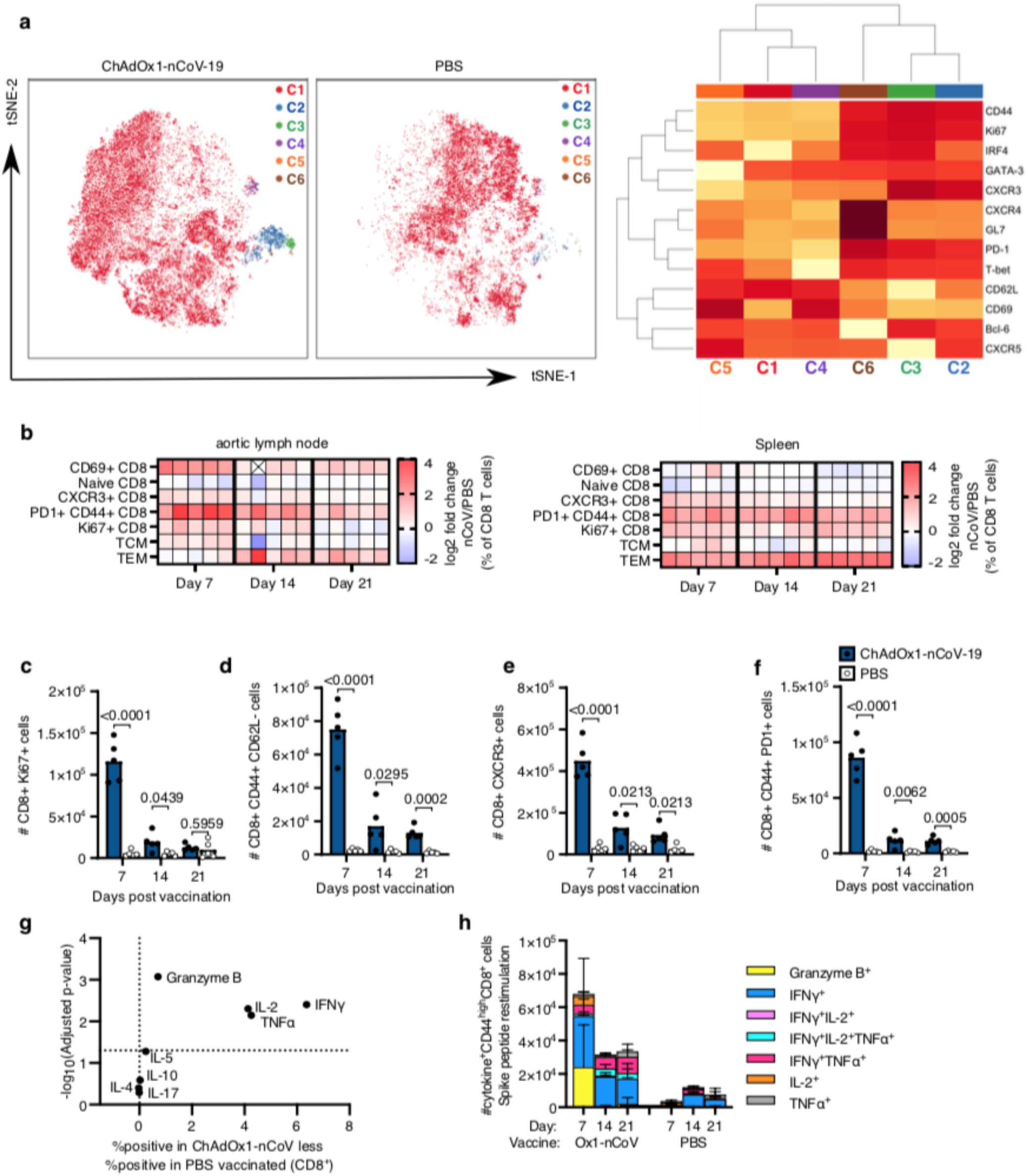
ChAdOx1 nCoV-19 induces a CD8 T cell response. **a.** tSNE/FlowSOM analyses of CD8^+^ T cells from 3-month-old (3mo) mice seven days after immunization with ChAdOx1 nCoV-19 or PBS. **b**. Heatmap of the manually gated CD8 T cell populations indicated at seven, 14 and 21 days after immunisation in the aortic lymph node and spleen. The frequency of each cell subset in each ChAdOx1 nCoV-19 immunised mouse has been expressed as the log2 fold change over the average frequency in PBS immunised mice (n=5). Crossed boxes indicate that there were none of that cell type for that mouse. Bar charts showing the number of Ki67^+^ (**c**), effector memory CD44^+^CD62L^-^ (**d**), CXCR3^+^ (**e**) and PD-1^+^CD44^+^ (**f**) CD8 cells in the aortic lymph node of ChAdOx1 nCoV-19 or PBS immunised mice, at the indicated timepoints post immunisation. **g**. Analysis of cytokine production six hours after PdBu/ionomycin stimulation of aortic lymph node cells from 3-month-old mice seven days after immunization with ChAdOx1 nCoV-19 or PBS. **h**. Stacked bar plots show the number of CD8^+^ cells singly or co-producing Granzyme B, IFNγ, IL-2 or TNFα six hours after restimulation with SARS-CoV-2 peptide pools, each bar segment represents the mean and the error bars the standard deviation. In c-f bar height in corresponds to the mean and each circle represents one biological replicate. P-values are calculated using a student’s t-test with Holm-Sidak multiple testing correction.

### A prime-boost strategy corrects dysregulated CD8 T cell priming in aged mice

To assess the CD8^+^ T cell response to ChAdOx1 nCoV-19 immunisation in the context of ageing, we immunised 3-month-old and 22-month-old mice and enumerated the CD8^+^ T cell types altered by vaccination (in Fig. 4) nine days after immunisation (**Fig. 5a**). In the draining aLN, CD8^+^ T cells from aged mice expressed markers of activation and proliferation in response to ChAdOx1 nCoV-19. Unlike in younger adult mice, the frequency of CXCR3^+^ cells or T effector memory cells did not increase in aged mice, compared to the PBS vaccinated group (**Fig. 5b-d**). At this early timepoint, the frequency of central memory T cells was not altered in either younger adult or aged mice by ChAdOx1 nCoV-19 vaccination (**Fig. 5e**). In the spleen, fewer Ki67^+^ CD8^+^ T cells were observed in aged mice after ChAdOx1 nCoV-19 vaccination, compared to younger adult mice (**Fig. 5f**). The formation of antigen-specific CD8^+^ T cells was assessed by restimulating splenocytes with SARS-CoV-2 spike protein peptide pools. Aged mice had a near absence of granzyme B producing CD8^+^ T cells, but production of IFNγ and TNFα was not significantly impaired compared to younger mice (**Fig. 5g**). IL-2 production was low in both adult and aged mice at this time point (**Fig. 5g**), compared to the seven days after immunisation (Fig. 4g-h). Despite a trend to lower cytokine production by CD8^+^ T cells in aged mice, the proportion of polyfunctional CD8^+^ T cells was not significantly diminished in aged mice after ChAdOx1 nCoV-19 vaccination (**Fig. 5h**). This demonstrates that a single dose of ChAdOx1 nCoV-19 induces an altered CD8^+^ T cell response in aged mice characterised primarily by a failure to form granzyme B-producing effector cells.

**Figure 5.**
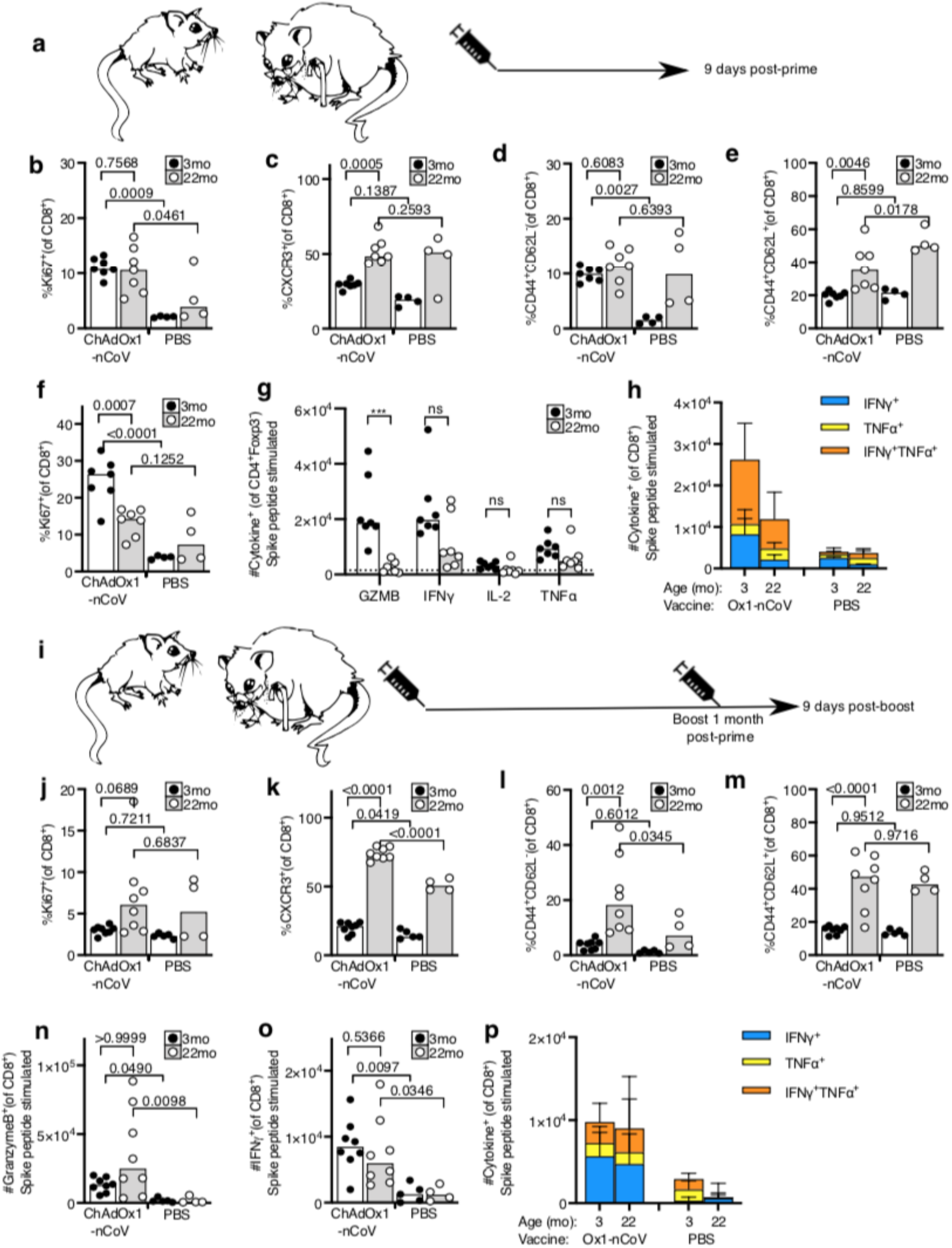
A prime-boost strategy enhances the CD8 T cell response to ChAdOx1 nCoV-19 in aged mice. **a**. Cartoon of prime immunization strategy. Percentage of Ki67^+^ (**b**), CXCR3^+^ (**c**), effector memory CD44^+^CD62L^-^ (**d**) and central memory CD44^+^CD62L^+^ (**e**) CD8^+^ T cells in the draining aortic lymph node from 3-month-old (3mo) or 22-month-old (22mo) mice nine days after immunization with ChAdOx1 nCoV-19 or PBS. **f**. Percentage of proliferating Ki67^+^ splenic CD8^+^ T cells in 3-month-old (3mo) or 22-month-old (22mo) mice nine days after immunization with ChAdOx1 nCoV-19 or PBS. **g-h**. Number of CD8^+^ T cells producing granzyme B (GZMB), IFNγ, IL-2 or TNFα six hours after restimulation with SARS-CoV-2 peptide pools, in (**g**) and the number of single and double cytokine producing CD8^+^ T cells are represented in stacked bar charts. Spleen cells are taken from 3-month-old (3mo) or 22-month-old (22mo) mice nine days after immunization with ChAdOx1 nCoV-19 or PBS. **i.** Cartoon of prime-boost immunization strategy. Percentage of Ki67^+^ (**j**), CXCR3^+^ (**k**), effector memory CD44^+^CD62L^-^ (**l**) and central memory CD44^+^CD62L^+^ (**m**) CD8^+^ T cells in the draining aortic lymph node from 3-month-old or 22-month-old mice nine days after immunization with ChAdOx1 nCoV-19 or PBS. **n**. Percentage of proliferating Ki67^+^ splenic CD8+ T cells in 3-month-old or 22-month-old mice nine days after immunization with ChAdOx1 nCoV-19 or PBS. Number of CD8^+^ cells producing Granzyme B (**n**) or IFNγ (**o**) six hours after restimulation with SARS-CoV-2 peptide pools, in (**p**) and the number of single and double cytokine producing CD8^+^ T cells are represented in stacked bar charts. Spleen cells are taken from 3-month-old or 22-month-old mice nine days after immunization with ChAdOx1 nCoV-19 or PBS. Bar height in **b-g, j-o** corresponds to the median and each circle represents one biological replicate. In **h, p**, each bar segment represents the mean and the error bars the standard deviation. The Shapiro-Wilk normality test was used to determine whether the data are consistent with a normal distribution, followed by either an ordinary one-way ANOVA test for data with a normal distribution or a Kruskal Wallis test for non-normally distributed data alongside a multiple comparisons test. Data are representative of two independent experiments (n=4-8 per group/experiment).

To determine whether a second dose could improve this response, we administered a booster dose of ChAdOx1 nCoV-19 one month after prime immunisation (**Fig. 5i**). Nine days after boost, an increase in Ki67^+^ CD4 T cells was not observed in the draining aLN (**Fig. 5j**), possibly due to the kinetics of the secondary response being faster than the primary. A significant increase in CXCR3^+^ CD8^+^ T cells and effector memory cells was observed in aged mice after boost, with no change in the proportion of central memory cells in either age group (**Fig. 5k-m**). Assessment of antigen-specific splenocytes showed that the booster dose of ChAdOx1 nCoV-19 rescued the production of granzyme B producing CD8^+^ T cells in aged mice (**Fig. 5n**). IFNγ production and cytokine polyfunctionality were similar to that following prime immunisation (**Fig. 5o, p**). This demonstrates that ChAdOx1 nCoV-19 is immunogenic in aged mice, and a booster dose can correct the age-dependent defect in the formation of granzyme B-producing CD8^+^ T cells.

### Prime-boost enhances the CD4^+^ T cell response to ChAdOx1 nCoV-19 in aged mice

Nine days after primary immunisation of aged mice (**Fig. 6a**), an increase in Ki67^+^CD4^+^ T cells and CXCR3-expressing Th1 cells was observed in the draining lymph node of ChAdOx1 nCoV-19 immunised mice compared to PBS immunised control mice (**Fig. 6b, c**). This was accompanied by an increase in the frequency of Th1-like Tregs in both adult and aged mice (**Fig. 6d**). An increased frequency of these cell types was likewise observed in the spleen in response to ChAdOx1 nCoV-19 immunisation in both adult and aged mice (**Fig. 6e-g**). It is notable that, by these measurements, the response in aged mice is comparable to that in younger adults. The antigen-specific CD4^+^ T cell response was assessed by restimulating splenocytes with SARS-CoV-2 spike protein peptide pools. As in young mice (Fig. 3), the response in aged mice to ChAdOx1 nCoV-19 was Th1 dominated, however there were fewer cytokine producing cells in aged mice nine days after a single immunisation (p<0.001, **Fig 6h, i**).

**Figure 6.**
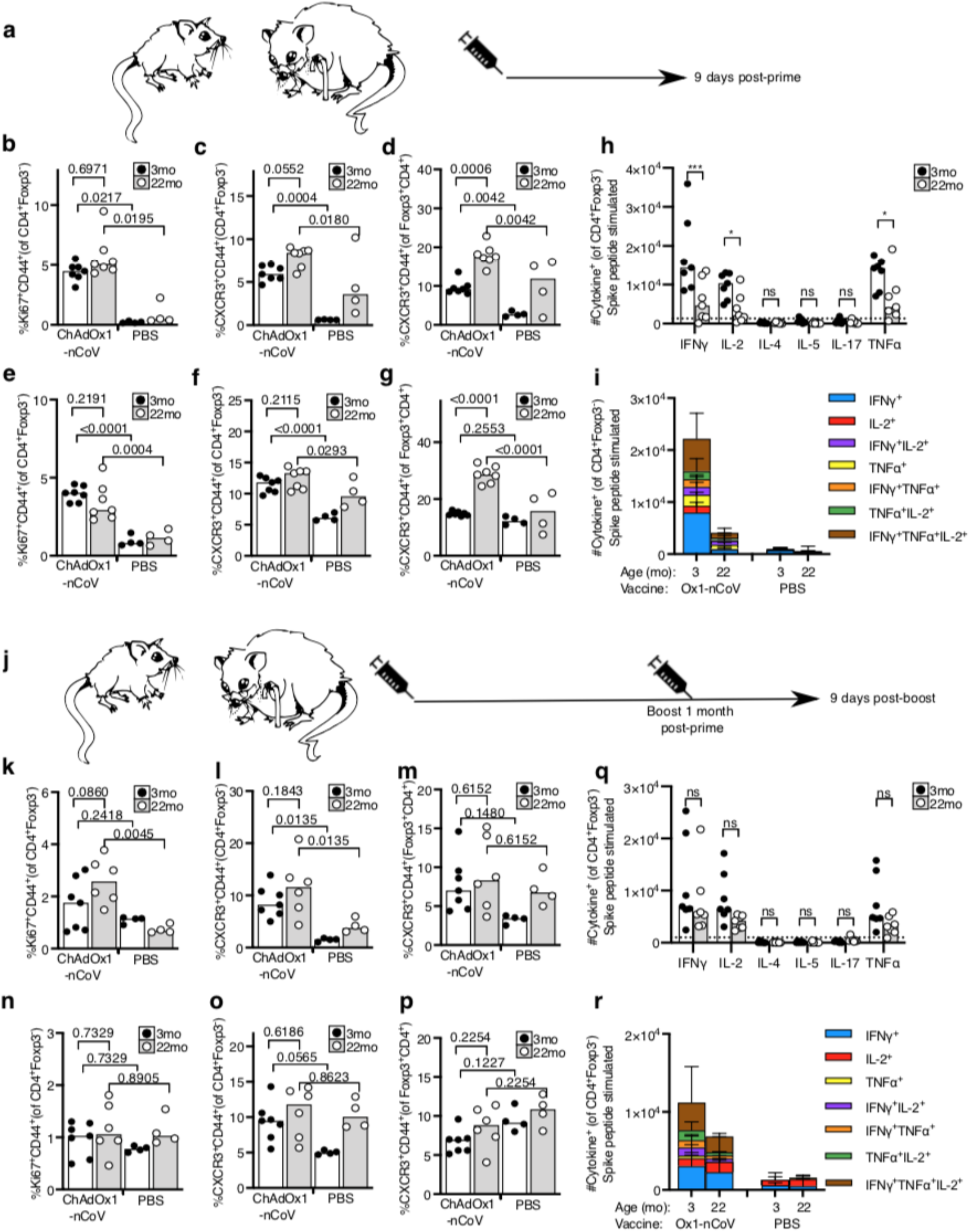
The CD4 cell response to ChAdOx1 nCoV-19 in aged mice. **a.** Cartoon of prime immunization strategy. Percentage of proliferating Ki67^+^ (**b**), CXCR3^+^CD44^+^ CD4 T cells (**c**) and CXCR3^+^CD44^+^Foxp3^+^ Treg cells (**d**) in the draining aortic lymph node. Percentage of proliferating Ki67^+^ (**e**), CXCR3^+^CD44^+^ CD4 T cells (**f**) and CXCR3^+^CD44^+^Foxp3^+^ Treg cells (**g**) in the spleen of 3-month-old or 22-month-old mice nine days after immunization with ChAdOx1 nCoV-19 or PBS. **h, i**. Number of CD4^+^Foxp3^-^ cells producing IFNγ, IL-2, IL-4, IL-5, IL-17 or TNFα six hours after restimulation with SARS-CoV-2 peptide pools, in (**i**) and the number of single and multiple cytokine producing CD4^+^ T cells are represented in stacked bar charts. **j.** Cartoon of prime-boost immunization strategy. Percentage of proliferating Ki67^+^ (**k**), CXCR3^+^CD44^+^ CD4 T cells (**l**) and CXCR3^+^CD44^+^Foxp3^+^ Treg cells (**m**) in the draining aortic lymph node. Percentage of Ki67^+^ CD44^+^ (**n**), CXCR3^+^CD44^+^ CD4^+^Foxp3^-^ T cells (**o**) and CXCR3^+^CD44^+^Foxp3^+^ Treg cells (**p**) in the spleen of 3-month-old or 22-month-old mice nine days after immunization with ChAdOx1 nCoV-19 or PBS. **q, r**. Number of CD4^+^Foxp3^-^ T cells producing IFNγ, IL-2, IL-4, IL-5, IL-17 or TNFα six hours after restimulation with SARS-CoV-2 peptide pools, in (**r**) and the number of single and multiple cytokine producing CD4^+^ T cells are represented in stacked bar charts. Bar height in corresponds to the median and each circle represents one biological replicate. In **i, r**, each bar segment represents the mean and the error bars the standard deviation. The Shapiro-Wilk normality test was used to determine whether the data are consistent with a normal distribution, followed by either an ordinary one-way ANOVA test for data with a normal distribution or a Kruskal Wallis test for non-normally distributed data alongside a multiple comparisons test. Data are representative of two independent experiments (n=4-8 per group/experiment).

A booster dose of ChAdOx1 nCoV-19 administered one month after prime (**Fig. 6j**) stimulated Ki67 expression and the formation of CXCR3^+^CD44^+^ Th1 cells, but not CXCR3^+^ Th1-like Treg cells in the draining lymph node of aged mice (**Fig. 6k-m**). In the spleen, the booster dose did not enhance the frequency of Ki67^+^ CD4^+^ T cells or the formation of CXCR3^+^ conventional or regulatory T cells in adult or aged mice (**Fig. 6n-p**). In contrast to the response to primary immunisation, the number of antigenspecific cytokine producing cells was comparable in adult and aged mice after booster immunisation (**Fig. 6q, r**). Together, this indicates that the CD4^+^ T cell response to ChAdOx1 nCoV-19 immunisation is largely intact in aged mice, with a slight deficiency in antigen-specific cytokine production that can be enhanced by a booster immunisation.

### Aged mice have an impaired germinal centre response after primary immunisation

The majority of clinically available vaccines are thought to provide protection by eliciting humoral immunity. Therefore it was important to quantify the B cell response to ChAdOx1 nCoV-19 vaccination in the context of ageing. Early antibody production after vaccination arises from antibody-secreting cells generated in the extrafollicular plasma cell response, which is fast, but typically short-lived(*38*). A comparable early plasma cell response was detected in the aLN of younger adult and aged mice after immunisation (**Fig. 7a, b**), although there was an increase in the proportion of IgM^+^ plasma cells in aged mice (**Fig. 7c**). An intact plasma cell response was coupled with an increase in serum antibodies nine days after immunization. These were of only slightly lower titre in aged mice, and of similar IgG subclass distribution to younger animals, indicative of a predominantly Th1 dominated response (**Fig. 7d-f**).

**Figure 7.**
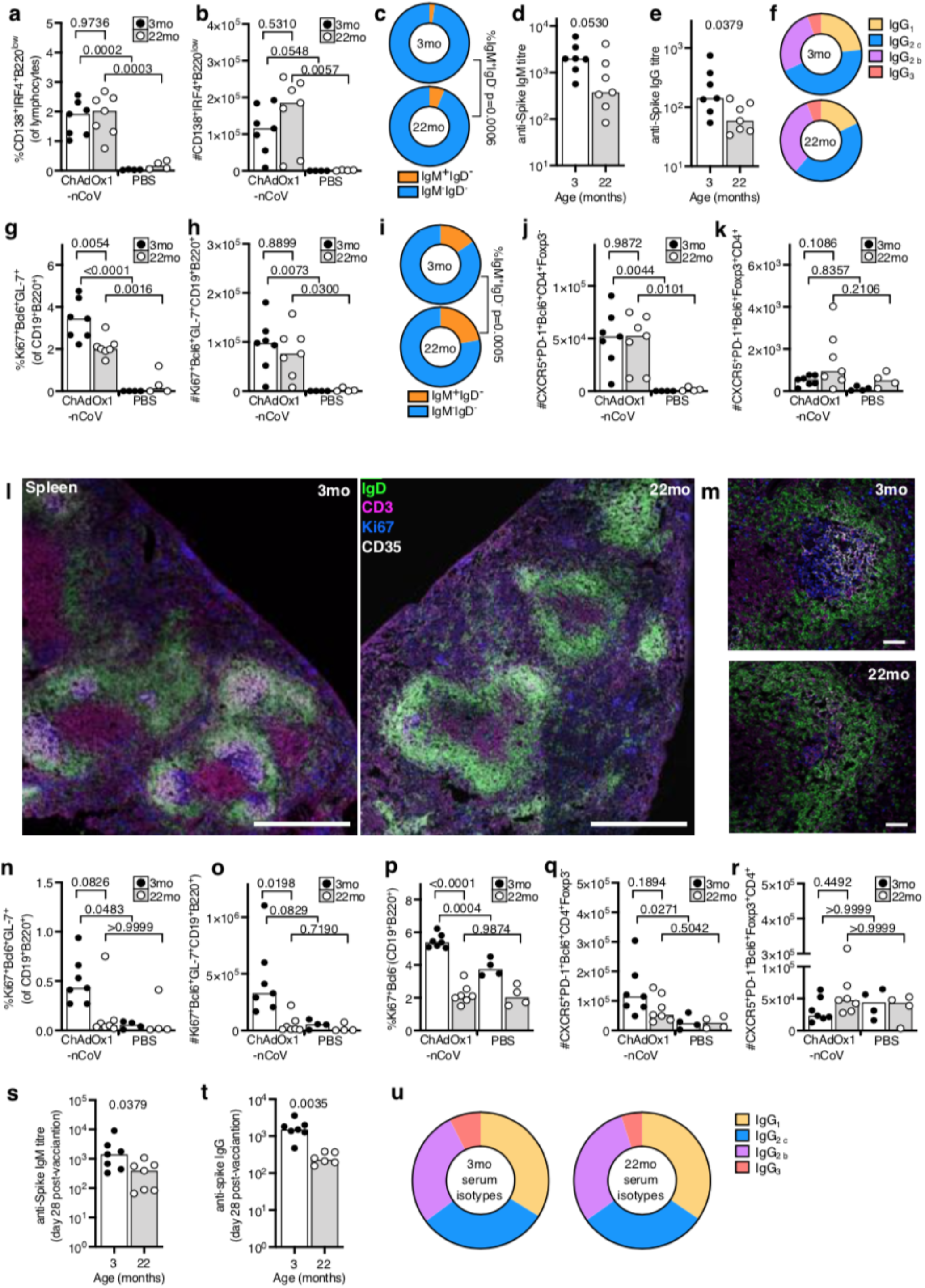
Impaired B cell responses after ChAdOx1 nCoV-19 immunisation of aged mice. B cell response in 3-month-old (3mo) or 22-month-old (22mo) mice nine days after immunization with ChAdOx1 nCoV-19 or PBS. Flow cytometric evaluation of the percentage (**a**) and number (**b**) of plasma cells in the aortic lymph node. **c**. Pie charts showing the proportion of IgM^+^IgD^-^ (orange) and switched IgM^-^IgD^-^ (blue) plasma cells from **b, c.** Serum IgM (**d**) and IgG (**e**) anti-spike antibodies nine days after immunization. **f.** Pie charts showing the proportion of anti-spike IgG of the indicated subclasses in the serum nine days after immunisation. Percentage (**g**) and number (**h**) of germinal centre B cells in the aortic lymph node. **i**. Pie charts showing the proportion of IgM^+^IgD^-^ (orange) and switched IgM^-^IgD^-^ (blue) germinal centre cells from **g, h**. Number of T follicular helper (**j**) and T follicular regulatory (**k**) cells in the draining lymph node. Confocal images of the spleen of ChAdOx1 nCoV-19 immunised mice of the indicated ages, in (**l**) the scale bars represent 500μm in (**m**) the scale bars represent 50μm. IgD^+^ B cell follicle in green, CD3^+^ T cells in magenta, Ki67^+^ cells in blue and CD35^+^ follicular dendritic cells in white. Percentage (**n**) and number (**o**) of splenic germinal centre B cells. **p**. Percentage of Ki67^+^ B cells in the spleen. Number of splenic T follicular helper (**q**) and T follicular regulatory (**r**) cells. Serum IgM (**e**) and IgG (**f**) anti-spike antibodies and IgG subclasses (**u**) 28 days after immunization. Bar height in corresponds to the median and each circle represents one biological replicate. The Shapiro-Wilk normality test was used to determine whether the data are consistent with a normal distribution, followed by either an ordinary one-way ANOVA test for data with a normal distribution or a Kruskal Wallis test for non-normally distributed data alongside a multiple comparisons test, for ELISA data analyses were done on log transformed values. Data are representative of two independent experiments (n=4-8 per group/experiment).

Long-lived antibody-secreting cells typically arise from the germinal centre response(*39*). The percentage, but not total number, of germinal centre B cells was reduced in aged mice compared to younger adult mice after ChAdOx1 nCoV-19 vaccination (**Fig. 7g, h**). Like the plasma cell response, there were more IgM^+^ germinal centre B cells in aged mice (**Fig. 7i**). An increase in T follicular helper cells, but not T follicular regulatory cells, accompanied the lymph node germinal centre response in adult and aged mice (**Fig. 7j, k**). In the spleen, germinal centres were easily visualised by microscopy in adult mice nine days after ChAdOx1 nCoV-19 vaccination, but were conspicuously absent in aged mice (**Fig. 7l, and Sup. Fig. 5**). Quantification of splenic germinal centres by flow cytometry confirmed impaired germinal centre formation in aged mice (**Fig. 7n, o**). This was accompanied by fewer proliferating non-germinal centre B cells and T follicular helper cells in aged mice (**Fig. 7p, q**). As in the draining lymph node, splenic T follicular regulatory cells were not induced by ChAdOx1 nCoV-19 vaccination at this time point (**Fig. 7r**). The impact of an impaired germinal centre response on vaccine-specific antibodies was observed 28 days after immunisation, with aged mice having lower titres of anti-spike IgM and IgG (**Fig. 7s, t**), but a similar profile of IgG subclasses (**Fig. 7u**). Together, these data indicate that whilst a single of dose of ChAdOx1 nCoV-19 can induce comparable extrafollicular plasma cell responses between young and aged mice, the germinal centre response is compromised with age.

### A second dose of ChAdOx1 nCoV-19 boosts humoral immunity in aged mice

To test whether a prime-boost strategy can enhance the B cell response in aged mice, a prime-boost approach was taken (**Fig. 8a**). Nine days after boost, there were Ki67^+^ non-germinal centre B cells, plasma cells and germinal centre B cells in the draining lymph nodes of aged mice (**Fig. 8b-h**). Notably, the magnitude of the germinal centre response was larger in aged mice than in younger adult mice after boost (**Fig. 8f-h**) and this was associated with increased T follicular helper and T follicular regulatory cell numbers (**Fig. 8i, j**). A germinal centre response was not observed in the spleen of either adult or aged mice nine days after booster immunisation (**Fig. 8k**). This demonstrates that a second dose of ChAdOx1 nCoV-19 can enhance the B cell response in aged mice. This improvement in the B cell response corresponded to an increase in anti-spike IgG, without skewing IgG subclasses, antibodies in every aged mouse that was given a booster immunisation (**Fig. 8l-o**). The post boost ratio of IgG_2_/IgG_1_ was 3.7 (3.0) in younger adult and 2.6 (1.6) in aged mice (mean and standard deviation). The functional effect of the humoral immunity after both prime and boost immunisations was measured by SARS-CoV-2 pseudotyped virus microneutralisation assay. Nine days after prime immunisation, SARS-CoV-2 neutralising antibodies were at a lower titre in aged mice than measured in adult mice (**Fig 8p**). Nine days after boost, neutralising antibodies were detectable in all aged mice and had been boosted eight-fold compared to post-prime, although the titre was significantly lower than in younger adult mice (**Fig 8q**). This demonstrates that a booster dose of ChAdOx1 nCoV-19 can improve vaccine-induced humoral immunity in older mice.

**Figure 8.**
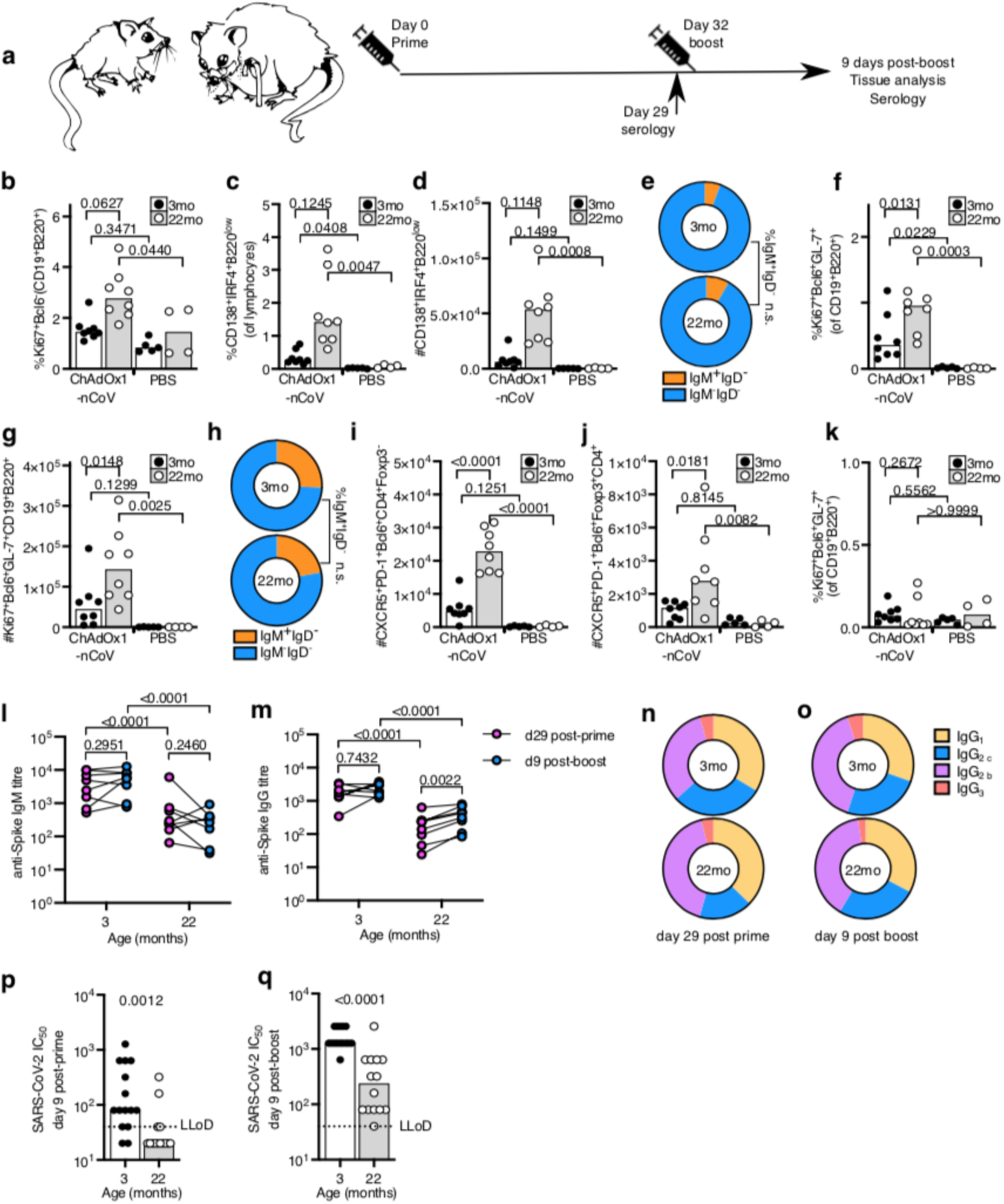
A booster immunization enhances the B cell response to ChAdOx1 nCoV-19 immunisation in aged mice. **a**. Scheme of the prime-boost immunization protocol. **b**. Percentage of Ki67^+^ B cells in the draining lymph node. Percentage (**c**) and number (**d**) plasma cells in the aortic lymph node. **e**. Pie charts showing the proportion of IgM^+^IgD^-^ (orange) and switched IgM^-^IgD^-^ (blue) plasma cells from b, c. Percentage (**f**) and number (**g**) of germinal centre B cells in the aortic lymph node. **h**. Pie charts showing the proportion of IgM^+^IgD^-^ (orange) and switched IgM^-^IgD^-^ (blue) germinal centre cells from g, h. Number of T follicular helper (**i**) and T follicular regulatory (**j**) cells in the draining lymph node. Percentage (**k**) of splenic germinal centre B cells. Serum anti-spike IgM (**l**), IgG (**m**) and IgG subclasses (**n, o**) prior to boost (day 29) and nine days after boost immunization. **p-q** SARS-CoV-2 neutralising antibody titres in sera were determined by micro neutralisation test, expressed as reciprocal serum dilution to inhibit pseudotyped virus entry by 50% (IC_50_). Samples below the lower limit of detection (LLoD) are shown as half of the LLoD. Bar height in corresponds to the median and each circle represents one biological replicate. The Shapiro-Wilk normality test was used to determine whether the data are consistent with a normal distribution, followed by either an ordinary one-way ANOVA test for data with a normal distribution or a Kruskal Wallis test for non-normally distributed data alongside a multiple comparisons test. For ELISA data analyses were done on log transformed values. In b-o, are shown from one of two independent experiments (n=4-8 per group/experiment), in p and q the data are pooled from two experiments.

## Discussion

The development of an effective anti-SARS-CoV-2 vaccine represents an opportunity to limit the health, social and economic consequences of the current pandemic. Since older adults are more likely to have severe health outcomes after infection, a vaccination strategy that provides protection for this group is particularly desirable. Here, we have used a pre-clinical model to determine the immunogenicity of ChAdOx1 nCoV-19 in older bodies. Whilst this vaccine stimulates an immune response in aged mice, it is of reduced magnitude compared to that seen in younger adult animals, consistent with previous immunisation studies(*21, 24, 27*). Importantly, the data presented here show that a second homologous immunisation at the same dose level was able to enhance the B cell, helper T cell and CD8^+^ T cell response in aged mice. In people, the same prime-boost approach had an acceptable safety profile and enhanced humoral immunity(*33*), indicating that this is a rational vaccination approach for use in the older members of our communities – arguably the sector of society most in need of an effective vaccine to prevent COVID-19.

After ChAdOx1 nCoV-19 immunisation of aged mice, a significant decrease in the proportion of class-switched germinal centre B cells and plasma cells was observed compared to young adult mice. This was linked with a lower titre of anti-spike antibodies after primary vaccination, with IgG being more affected than IgM by age. Previous studies, in both humans and mice, have demonstrated that ageing causes a B cell-intrinsic impairment in class-switch recombination. This is attributed to a decrease in the expression of the enzyme activation-induced cytidine deaminase, which initiates class-switch, and its positive regulator, the E2A-encoded E47 transcription factor, in B cells from aged mice and humans(*40, 41*). The reduced expression of activation-induced cytidine deaminase in B cells from older persons has been correlated with lower neutralising antibody production after seasonal influenza vaccination(*42*). Therefore, vaccination strategies that aim to enhance class-switch recombination may improve vaccine efficacy in older people.

The age-dependent defects in the B cell response were not limited to class-switch recombination. This highlights that either the formation of antibody secreting cells, or their ability to secrete antibody is impaired by ageing. Antibody-secreting cells can form from two cellular pathways, but these are not equal in terms of quality or longevity. The first pathway, the extrafollicular response, produces an initial burst of antibodies early after antigenic challenge. This response is short-lived with no additional diversification of the B cell repertoire and thus its contribution to long-term immunity is minimal(*38*). The second pathway, the germinal centre reaction, is a specialised microenvironment that produces memory B cells and long-lived antibody secreting plasma cells with somatically-mutated immunoglobulin genes(*39, 43, 44*). The germinal centre is the only cellular source of long-lived plasma cells(*45*), and the only site where the B cell response can be altered in response to antigen. Nine days after ChAdOx1 nCoV-19 immunisation, the formation of plasma cells was comparable in young and aged mice, a time at which the majority of plasma cells derive from the extrafollicular response(*45*). By contrast, the germinal centre response was diminished in aged mice, which is a well described deficit of the older immune system after vaccination(*21, 46*). This indicates that poor germinal centre formation is the major barrier that must be overcome to improve humoral immunity in older individuals. In this study we show that a booster dose of ChAdOx1 nCoV-19 enhanced the magnitude of the germinal centre response in aged mice, to a greater extent than it did in younger animals. This may be because younger mice have produced higher titres of anti-vaccine antibodies, which limits the action of a second dose. A similar phenomenon occurs with the seasonal influenza vaccine, in which individuals with high titres of vaccine reactive antibodies typically have a muted response to secondary vaccination(*47*), and has been observed in an adenovirus type-5(Ad5)-vectored COVID-19 vaccine in people who have high pre-existing antibody titres to the vaccine vector due to natural exposure to Ad5(*48*).

Another feature of the aged germinal centre reaction is reduced selection of high affinity B cells, resulting in a diminished quality of the response in ageing(*23*). Whether the further diversification and affinity-based selection of the B cell receptor is important for protective humoral immunity to SARS-CoV-2 has yet to be established. However, antibodies isolated from spike-specific B cells from convalescent COVID-19 patients can be both potently neutralising and have low levels of somatic hypermutation(*49–51*), indicating that extensive somatic hypermutation is not required for anti-SARS-CoV-2 neutralising antibody formation. Therefore, the main barrier to inducing protective humoral immunity to SARS-CoV-2 in older people may be an issue of enhancing the magnitude, rather than quality, of the germinal centre response. Our study shows that it is possible to enhance the number of germinal centre B cells in aged mice by giving a second dose of the vaccine.

It has been proposed that a failure of the germinal centre reaction in older mice is caused by an increased frequency of suppressive T follicular regulatory cells(*52*). Here, we did not see an accumulation of T follicular regulatory cells in ageing, and their formation was only robustly observed later in the response. This result is similar to the delayed formation of T follicular regulatory cells after influenza A infection(*53*). Because the germinal centre response requires the coordinated response of many cell types, identifying the causal defect is challenging. In heterochronic parabiosis experiments, the age of the microenvironment, and not the age of the recirculating lymphocytes, underpins the poor germinal centre response in ageing(*54*). This implicates non-migratory cells, such as stromal cells or tissue resident cells as causal factors in the diminished germinal centre reaction in ageing.

A rapid recall of memory CD8^+^ T cells in response to viral infections complements the humoral response by promoting efficient pathogen clearance and this becomes particularly important in scenarios where the protective ability or magnitude of neutralising antibodies is compromised(*55*). A single dose of ChAdOx1 nCoV-19 vaccine induces a CD8^+^ T cell response in both adult and aged mice. In aged mice there was a trend for fewer cytokine producing CD8^+^ T cells, but this was not statistically significant in either the discovery or replication cohort. There was, however, a profound defect in the generation of spike-reactive CD8^+^ T cells that produced granzyme B, a key cytotoxic effector molecule produced by effector CD8 ^+^ T cells. This failure to generate granzyme B^+^ CD8^+^ T cells is a common feature in ageing, previously observed in aged mice after West Nile Virus infection(*56*) and in aged monkeys infected with SARS-CoV(*57*). A similar trend for fewer granzyme B^+^ CD8 cells in older people was observed after influenza vaccination, although this study was not sufficiently powered for the difference to be statistically significant(*58*). Consistent with reduced expression of granzyme B in T cells from older adults, two weeks after influenza vaccination T cells taken from older adults have an impaired ability to kill influenza infected cells *ex vivo(59).* In a small study of older people, poor induction of granzyme B after influenza vaccination was associated with a greater likelihood of subsequent laboratory confirmed influenza infection, indicating that enhancing granzyme B^+^ CD8^+^ T cells may be important for protective immunity against respiratory viruses in older adults(*60*). Here we demonstrate that a second dose of ChAdOx1 nCoV-19 was able to correct the defective granzyme B^+^ CD8^+^ T cell response in aged mice, indicating that two doses of this vaccine is a better approach to enhance the cellular immune response in older bodies.

ChAdOx1 nCoV-19 is currently being trialed in older adults as part of a phase III trial, which will ultimately determine whether it is an effective vaccine for this demographic. Immunisation of people over 55 years of age with Ad5-vectored COVID-19 vaccine resulted in lower antibody titres in the older age group, suggesting that adenoviral vectored vaccine strategies may require more than one dose in older people(*48*). However, as pre-existing immunity to Ad5, a naturally occurring human tropic adenovirus, can inhibit the response to this vaccine, it is possible that the reduced immunogenicity of this vaccine is due to an increased prevalence of antibodies to the vaccine vector in this age group. By contrast, the presence of anti-ChAdOx1 antibodies in the general population is low(*61*). The work presented here demonstrates that one dose of this vaccine is immunogenic in aged mice, but this response can be significantly improved with a second booster dose. Given that a second dose of ChAdOx1 nCoV-19 is immunogenic with expected reactogenicity profile in humans(*33*), this may be a viable strategy to enhance immunogenicity and possibly efficacy in older people.

## Methods

### Mouse housing and husbandry

C57BL/6Babr mice were bred, aged and maintained in the Babraham Institute Biological Support Unit (BSU). No primary pathogens or additional agents listed in the FELASA recommendations(*62*) were detected during health monitoring surveys of the stock holding rooms. Ambient temperature was ~19-21°C and relative humidity 52%. Lighting was provided on a 12 hr light: 12 hr dark cycle including 15 min ‘dawn’ and ‘dusk’ periods of subdued lighting. After weaning, mice were transferred to individually ventilated cages with 1–5 mice per cage. Mice were fed CRM (P) VP diet (Special Diet Services) ad libitum and received seeds (e.g. sunflower, millet) at the time of cage-cleaning as part of their environmental enrichment. All mouse experimentation was approved by the Babraham Institute Animal Welfare and Ethical Review Body. Animal husbandry and experimentation complied with existing European Union and United Kingdom Home Office legislation and local standards (PPL: P4D4AF812). Young mice were 10–12 weeks old, and aged mice 93–96 weeks old when experiments were started. Mice that had tumours, which can occur in aged mice, were excluded from the analysis.

### Immunisation and tissue sampling

Mice were immunised in the right quadriceps femoris muscle with 50μL of either 1×10^8^ infectious units of ChAdOx1 nCoV-19 in phosphate buffered saline (PBS) PBS alone, or 0.02μm yellow-green fluorescent Carboxylate-Modified Microspheres (Invitrogen # F8787) in phosphate buffered saline. At the indicated timepoints post vaccination, blood, the right aortic lymph node, spleen and right quadriceps femoris muscle were taken for analysis.

### Flow cytometry

For T and B cell flow cytometric stains a single cell suspension was prepared from the aortic lymph node and half the spleen was generated by pressing the tissues through a 70 μm mesh in 2% FBS in PBS. Cell numbers and viability were determined using a CASY TT Cell Counter (Roche). 2×10^6^ cells were transferred to 96-well plates for antibody staining. Samples were blocked with 100μl of 2.4G2 Fc Block (made in house) for 20 min at 4°C. Cells were then stained with surface antibody mix for 2hrs at 4°C and then were fixed with the eBiosciences Foxp3/Transcription Factor Staining Buffer (#00-5323-00) for 30 min at 4°C. Cells were then washed with 1x Permeabilisation buffer (eBioscience #00-8333-56) twice and stained with intracellular antibody mix in 1x Permeabilisation buffer at 4°C overnight. For cytokine staining, splenic cells were stimulated with a pool of SARS-CoV-2 spike protein immunodominant domain peptides, (Miltenyi Biotec #130-126-700) at a 0.6 μM concentration (approx.1μg/ml), whilst lymph node cells were stimulated with 0.5μg/ml of Phorbol 12,13 dibutyrate (PdBu, Tocris Bioscience, #), 0.75μg/ml of Ionomycin calcium salt (Tocris Bioscience, #), both in warm complete RPMI (10% FCS, 1% Pen/Strep, 1% glutamine, 1% sodium pyruvate, 1% MEM NAA, 1% HEPES and 55μM-2-mercaptoethanol) for 4hrs at 37°C, 5%CO_2_. Cytokine secretion was then blocked with 22μg/ml of Brefeldin A (Tocris Bioscience, #) in warm complete RPMI for 2hrs at 37°C, 5%CO_2_. The cells were then stained with surface antibody mix for 20 minutes at 4°C and were subsequently fixed with 2% formaldehyde for 30min at room temperature. After two wash steps with 1x Permeabilisation buffer (eBioscience #00-8333-56), the cells were stained with intracellular antibody mix in 1x Permeabilisation buffer, supplemented with 20% 2.4G2 Fc Block at 4°C overnight. Following overnight staining, samples were washed twice with 1x Permeabilisation buffer and once with 2% FBS in PBS and acquired on a Cytek™ Aurora. Cells for single colour controls were prepared in the same manner as the fully stained samples. The antibodies used for surface and overnight staining are listed in Table 1.

For dendritic cell analysis, secondary lymphoid tissues were harvested and incubated with 10 mg/ml Collagenase D (Roche #11088866001) in plain RPMI medium (Gibco #11875093) for 30-45 min at 37 °C, followed by gentle pipetting to disrupt the tissue. Cells were washed with PBS containing 2% FBS, before cell numbers were determined using a CASY TT Cell Counter (Roche). 4×106 cells were transferred to 96-well plates for antibody staining. After a wash with PBS, cells were stained with Live/Dead Fixable Blue Dead Cell Stain (Invitrogen #L23105; diluted 1:1000 in PBS) on ice for 10min after which they were washed in 2% FBS in PBS and treated with unlabelled anti-CD16/32 (eBioscience #14-0161-82; diluted 1:100 in 2% FBS in PBS) for 20min at 4°C. Surface antibody staining was then performed for 1hr at 4°C in Brilliant Stain Buffer (BD Biosciences #563794). Samples were acquired on an LSRFortessa 5 with stained UltraComp eBeads Compensation Beads (Invitrogen #01-2222-41) as compensation controls.

Manual gating of flow cytometry data was done using FlowJo v10 software (Tree Star). tSNE, FlowSOM and heatmap analysis were performed on aLN samples from day 7 postvaccination using R (version 4.0.2) using code that has previously been described(*63*). The antibodies used for surface staining are listed in Table 1.

### Confocal imaging

For imaging of germinal centres, half of the spleen was fixed in periodate-lysine-paraformaldehyde (PLP) containing 1% (v/v) PFA (Sigma #P6148), 0.075 M L-Lysine (Sigma #L5501), 0.37 M Na3PO4 (pH 7.4) (Sigma #342483) and 0.01 M NaIO4 (Sigma #210048), for 4 hr at 4 °C. For imaging yellowgreen fluorescent Carboxylate-Modified Microspheres (Invitrogen # F8787), lymph nodes and spleen were fixed in BD Cytofix/Cytoperm (BD Cat #554722), diluted 1 in 4 in PBS. After fixation, all tissues were dehydrated in 30% sucrose (Sigma #S0389) overnight, embedded in Optimum Cutting Temperature (OCT) medium (VWR #25608–930) on dry ice and stored at −80 °C. The frozen tissues were cut into 10μm sections using a cryostat (Leica Biosystems) at −20°C and again stored at −80°C. For imaging yellow-green fluorescent carboxylate-codified microspheres, slides were first air-dried and then washed in PBS three times after which DAPI staining was performed (Invitrogen #D1306; diluted 1:10 000 in PBS) for 10 minutes at room temperature protected from light. Slides were then washed three times with PBS and coverslips were mounted on the slides using Hydromount mounting medium (National Diagnostics #HS-106). For antibody stains, the slides were first air-dried and then hydrated in 0.5% Tween 20 in PBS (PBS-T). The slides were blocked in PBS containing 2% BSA and 10% goat serum for 2hrs, washed three times with PBS-T and then permeabilised with PBS containing 2% Triton X (Sigma #X100) for 45min. Following three wash steps with PBS-T, the slides were stained with primary antibody mix, which included AF647-conjugated rat anti-mouse IgD (clone 11-26 c.2a, Biolegend; 1:200), FITC-conjugated rat anti-mouse Ki67 (clone SolA15, Invitrogen; 1:100), rat antimouse biotin-conjugated CD21/35 (clone 8D9, ThermoFisher Scientific; 1:400) and hamster antimouse CD3ε (clone 500A2, ThermoFisher Scientific; 1:200), in PBS-T containing 1% BSA at 4°C overnight. The next day, slides were washed three times with PBS-T and incubated with secondary antibody mix, which included AF568-conjugated goat anti-hamster IgG (Thermofisher, 1:1000) and BV421-conjugated Streptavidin (Biolegend, 1:1000), in PBS-T containing 2% goat serum for 2hr at room temperature. Slides were then washed three times with PBS-T, PBS and dH_2_O and coverslips were mounted using Hydromount mounting medium (National Diagnostics #HS-106). Images were acquired using a Zeiss 780 confocal microscope with 10x and 20x objectives and analysed using ImageJ.

### RNA isolation and quantitative Real-Time PCR

The right quadriceps muscle was dissected and weighed. The muscle tissue was snap frozen in liquid nitrogen and homogenized in 2mL TRIzol reagent (Thermo Fisher Scientific #15596026), then 1 mL of the homogenized muscle in TRIzol was processed following the manufacturer’s instructions. RNA concentrations obtained from the RNA isolation were measured using the NanoDrop system (Thermo Fisher Scientific). RT-qPCR was performed using TaqMan Gene Expression Assays for *Mx1* (Mm 00487796_m1), *Gbp2* (Mm 00494576_g1) and *Hprt* (Mm 03024075_m1) directly on RNA using Thermo Fisher Scientific’s TaqMan RNA-to-CT 1-Step Kit (#4392656) following the manufacturer’s protocol. All RT-qPCR reactions were assembled in 384-well plates (Bio-Rad #HSP3805), adding 10ng of template RNA per reaction to 8μl of a master mix containing the appropriate TaqMan Gene Expression Assay. All samples were run in triplicates on a BioRad CFX384 Real-Time System. The 2^-ΔΔCt^-method was applied for relative quantification of mRNA levels. Samples from PBS immunised mice were used as calibrators. Cq values were exported from the CFX Manager software (Bio-rad).

### Enzyme-linked immunosorbent assay

Standardised ELISA was performed to detect SARS-CoV-2 FL-S protein – specific antibodies in sera. MaxiSorp plates (Nunc) were coated with 100 or 250 ng/well FL-S protein overnight at 4°C for detection of IgG or IgM and IgA, respectively, prior to washing in PBS/Tween (0.05% v/v) and blocking with Blocker Casein in PBS (Thermo Fisher Scientific) for 1 h at room temperature (RT). Standard positive serum (pool of mouse serum with high endpoint titre against FL-S protein), individual mouse serum samples, negative and an internal control (diluted in casein) were incubated for 2h at RT for detection of specific IgG or 1h at 37°C for detection of specific IgM or IgA. Following washing, bound antibodies were detected by addition of AP-conjugated goat anti-mouse IgG (Sigma-Aldrich) for 1h at RT or addition of AP-conjugated goat anti-mouse IgM or IgA (Abcam and Sigma-Aldrich, respectively) and addition of pNPP substrate (Sigma-Aldrich). An arbitrary number of ELISA units were assigned to the reference pool and OD values of each dilution were fitted to a 4-parameter logistic curve using SOFTmax PRO software. ELISA units were calculated for each sample using the OD values of the sample and the parameters of the standard curve.

The IgG subclass ELISA were performed according to the protocol described for detection of specific IgM or IgA in the serum. In addition, all serum samples were diluted to 1 total IgG ELISA unit and then detected with anti-mouse IgG subclass-specific secondary antibodies (Southern Biotech or Abcam). The results of the IgG subclass ELISA are presented using OD values instead of the ELISA units used for the total IgG ELISA. The ratio of IgG_2_/IgG_1_ was calculated for each animal as sum of OD values (IgG_2b_ + IgG_2c_) divided by the OD value of IgG_1_ and represented as mean values with standard deviation (SD).

### Micro neutralisation test using lentiviral-based pseudotypes bearing the SARS-CoV-2 Spike

Lentiviral-based SARS-CoV-2 pseudotyped viruses were generated in HEK293T cells incubated at 37 °C, 5% CO_2_ as previously described(*32*). Briefly, cells were seeded at a density of 7.5 *x* 10^5^ in 6 well dishes, before being transfected with plasmids as follows: 500 ng of SARS-CoV-2 spike, 600 ng p8.91 (encoding for HIV-1 gag-pol), 600 ng CSFLW (lentivirus backbone expressing a firefly luciferase reporter gene), in Opti-MEM (Gibco) along with 10 μL PEI (1 μg/mL) transfection reagent. A ‘no glycoprotein’ control was also set up using the pcDNA3.1 vector instead of the SARS-CoV-2 S expressing plasmid. The following day, the transfection mix was replaced with 3 mL DMEM with 10% FBS (DMEM-10%) and incubated for 48 and 72 hours, after which supernatants containing pseudotyped SARS-CoV-2 (SARS-CoV-2 pps) were harvested, pooled and centrifuged at 1,300 x *g* for 10 minutes at 4 °C to remove cellular debris. Target HEK293T cells, previously transfected with 500 ng of a human ACE2 expression plasmid (Addgene, Cambridge, MA, USA) were seeded at a density of 2 × 10^4^ in 100 μL DMEM-10% in a white flat-bottomed 96-well plate one day prior to harvesting SARS-CoV-2 pps. The following day, SARS-CoV-2 pps were titrated 10-fold on target cells, and the remainder stored at −80 °C. For micro neutralisation tests, mouse sera were diluted 1:20 in serum-free media and 50 μL was added to a 96-well plate in triplicate and titrated 2-fold. A fixed titred volume of SARS-CoV-2 pps was added at a dilution equivalent to 10^5^ signal luciferase units in 50 μL DMEM-10% and incubated with sera for 1 hour at 37 °C, 5% CO_2_ (giving a final sera dilution of 1:40). Target cells expressing human ACE2 were then added at a density of 2 *x* 10^4^ in 100 μL and incubated at 37 °C, 5% CO_2_ for 72 hours. Firefly luciferase activity was then measured with BrightGlo luciferase reagent and a Glomax-Multi^+^ Detection System (Promega, Southampton, UK). Pseudovirus neutralization titres were expressed as the reciprocal of the serum dilution that inhibited luciferase expression by 50% (IC_50_).

### Statistics

All experiments were performed either twice or three times with 3–8 mice per group. Data was first tested for gaussian distribution using a Shapiro-Wilk test. Then data that was consistent with a normal distribution was analysed with either a student’s t-test for comparing two data set, or one-way ANOVA test for data with multiple groups. If the data did not follow a normal distribution then a Mann-Whitney test was used for comparing two data sets and a Kruskal Wallis test for multiple comparisons. For ELISA data p-values were generated on log transformed data. All p-values shown are adjusted for multiple comparisons where multiple tests were performed on the same data. Analyses were performed within the Prism v8 software (GraphPad).

## Acknowledgements

We are grateful to Dr. Martin Turner, Dr. Geoff Butcher, Marc Wiltshire, Paul Symonds and Prof. Michael Wakelam for their support. This study was supported by funding from the Biotechnology and Biological Sciences Research Council (BBS/E/B/000C0427, BBS/E/B/000C0428 and the Campus Capability Core Grant to the Babraham Institute), the Lister institute of Preventative Medicine, the EPSRC VaxHub (EP/RO13756/1) and Innovate UK (biEBOV: 971615). Teresa Lambe and Sarah Gilbert are Jenner Investigators. Michelle Linterman is an EMBO Young Investigator and a Lister Institute Prize Fellow. Jia Le Lee is supported by a National Science Scholarship (PhD) by the Agency for Science, Technology and Research, Singapore.

## Author contributions

Conceptualisation, M.A.L. and T.L.; Methodology, A.S.C., W.S.F., S.I., O.T.B., A.J.S., S. B-R., D.W., J.B., D.B., S.F-B. M.A.L., N.T., C.C., N. E-B., and C.N.; Investigation, A.S.C., W.S.F., S.I., O.T.B., S.F-B., A.J.S., N.T., C.C., S. B-R., J.L.L., M.A.L; Writing—Original Draft Preparation, M.A.L., A.S.C., W.S.F., J.L.L., and T.L.; Writing—Review and Editing, All authors; Project Administration, S.I., M.A.L, and T.L.; Funding Acquisition, M.A.L, S.C.G, and T.L.

---

**Supplementary Figure 1.**
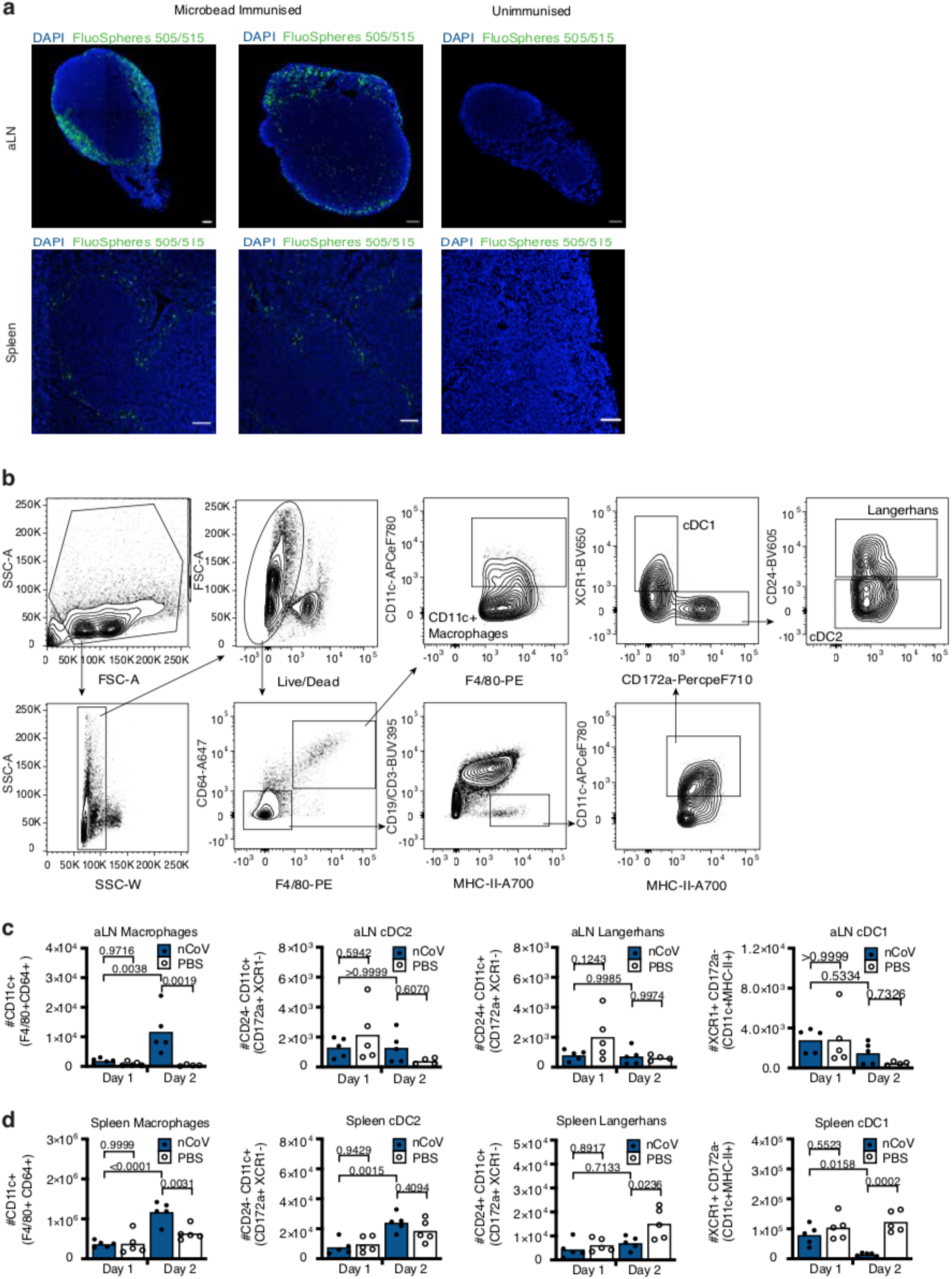
Intramuscular immunisation of FluoSpheres drains to the aortic LN and spleen where vaccination with ChAdOx1 nCoV results in the expansion of CD11c^+^ macrophages. **a.** Additional representative immunofluorescence confocal images of DAPI expression and FluoSpheres™ (505/515) localisation in the aLN (top) and spleen (bottom) of mice immunised with yellow-green fluorescent FluoSpheres™ (left and middle) or PBS (right) at 24hr post i.m immunisation. Scale bars, 100μm for both aLN and spleen. Data are representative of two independent experiments (n=2-5 per group/experiment). **b.** Gating strategy used for the identification of antigen presenting cell populations the aLN and spleen of mice immunised with ChAdOx1 nCoV-19 or PBS at one- and two-days post immunisation. Macrophages were defined as CD11c^+^F4/80^+^CD64^+^ cells. cDC1s were defined as XCR1^+^CD172a^-^CD11c^+^MHC-II^+^CD3/CD19^-^CD64^-^F4/80^-^ cells. cDC2s CD24^-^ CD172a^+^XCR1^-^CD11c^+^MHC-II^+^CD3/CD19^-^CD64^-^F4/80^-^ cells. Langerhans cells were defined as CD24^+^CD11 c^+^CD172a^+^XCR1^-^CD11 c^+^MHC-II^+^CD3/CD19^-^CD64^-^F4/80^-^ cells. **c-d.** Quantification of the absolute number of macrophages, cDC2s, Langerhans cells and cDC1s in the aLN (**c**) and spleen (**d**.) Bar height in **c-d** corresponds to the mean and each circle represents one biological replicate. P-values were determined using a Shapiro-Wilk normality test followed by either an ordinary one-way ANOVA test for data with a normal distribution or a Kruskal Wallis test for non-normally distributed data alongside a multiple comparisons test. Data are representative of two independent experiments (n=4-5 per group/experiment).

**Supplementary Figure 2.**
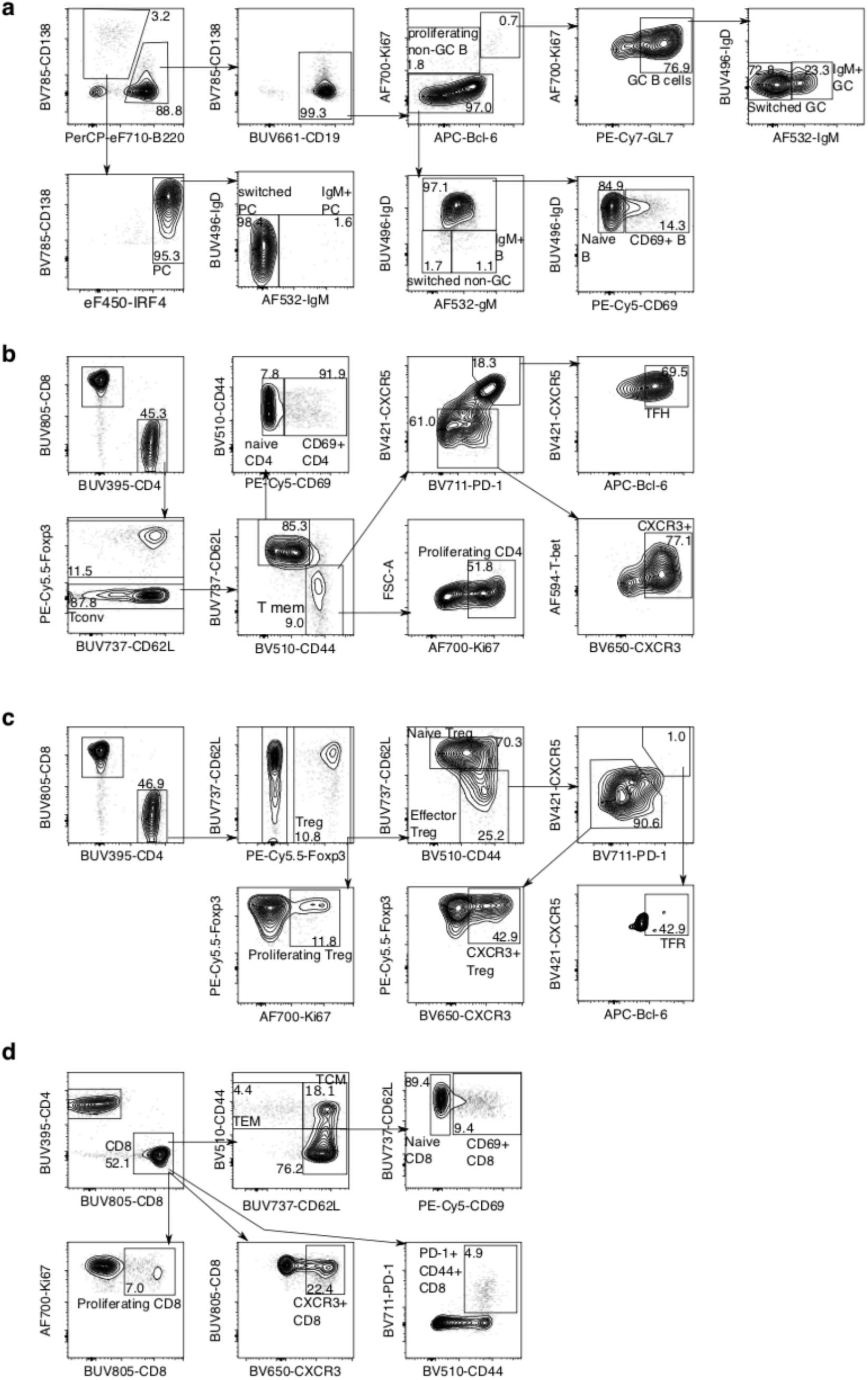
Gating strategy for ex vivo lymphocyte analyses. Representative flow cytometric contour plots showing the gating strategy for B cell subsets (**a**), CD4^+^Foxp3^-^ T cell subsets (**b**), Treg cell subsets (**c**) and CD8 cell subsets (**d**). Samples are taken from an aortic lymph node seven days after ChAdOx1 nCoV-19 immunisation.

**Supplementary Figure 3.**
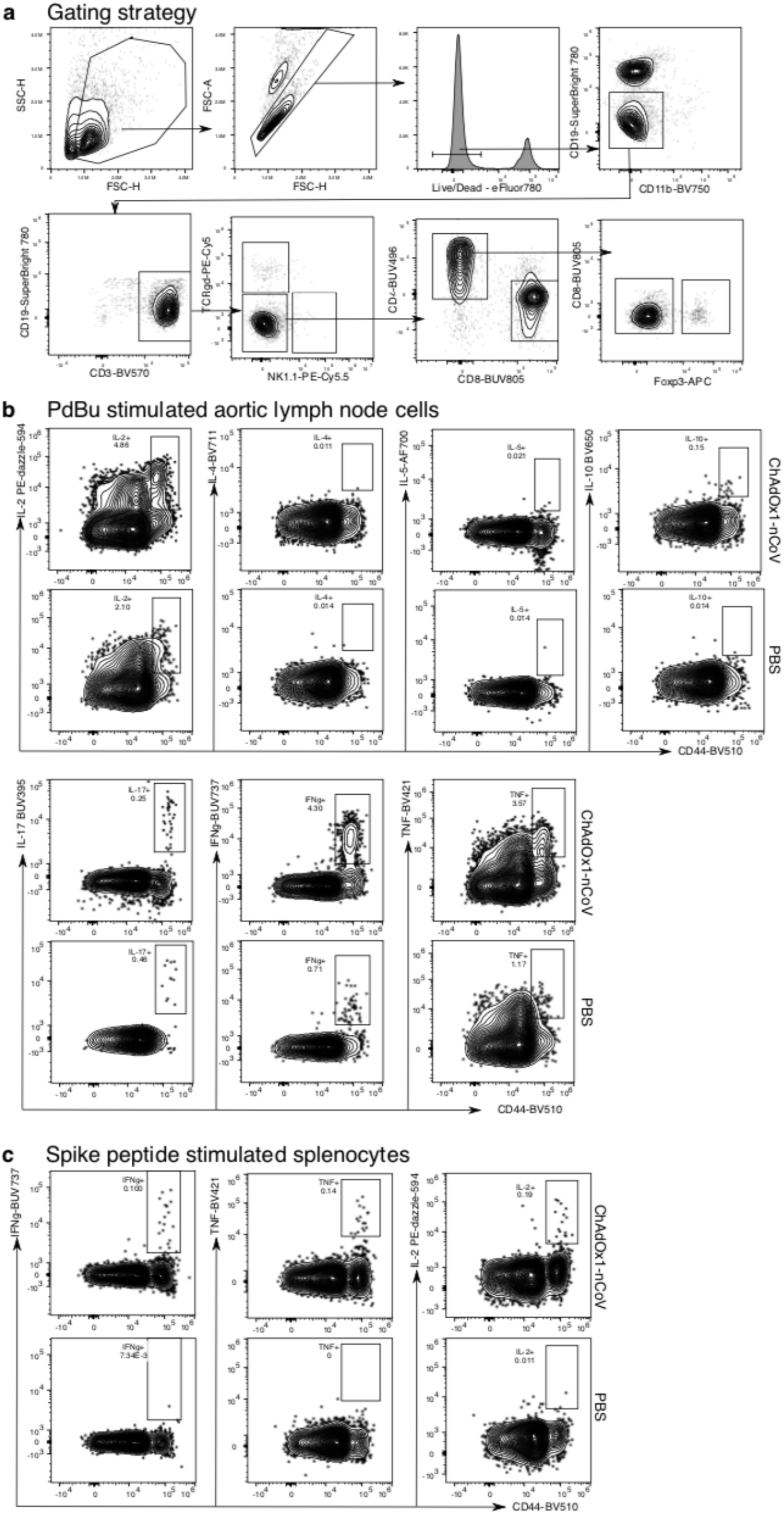
Gating strategy CD4 T cell cytokine staining. Flow cytometric contour plots showing the gating strategy to identify T cell subsets (**a**), cytokine production in CD4^+^Foxp3^-^ aortic lymph node T cells six hours after PdBu/Ionomycin restimulation (**b**), and in SARS-CoV-2 peptide restimulated CD4^+^Foxp3^-^ splenocytes (**c**).

**Supplementary Figure 4.**
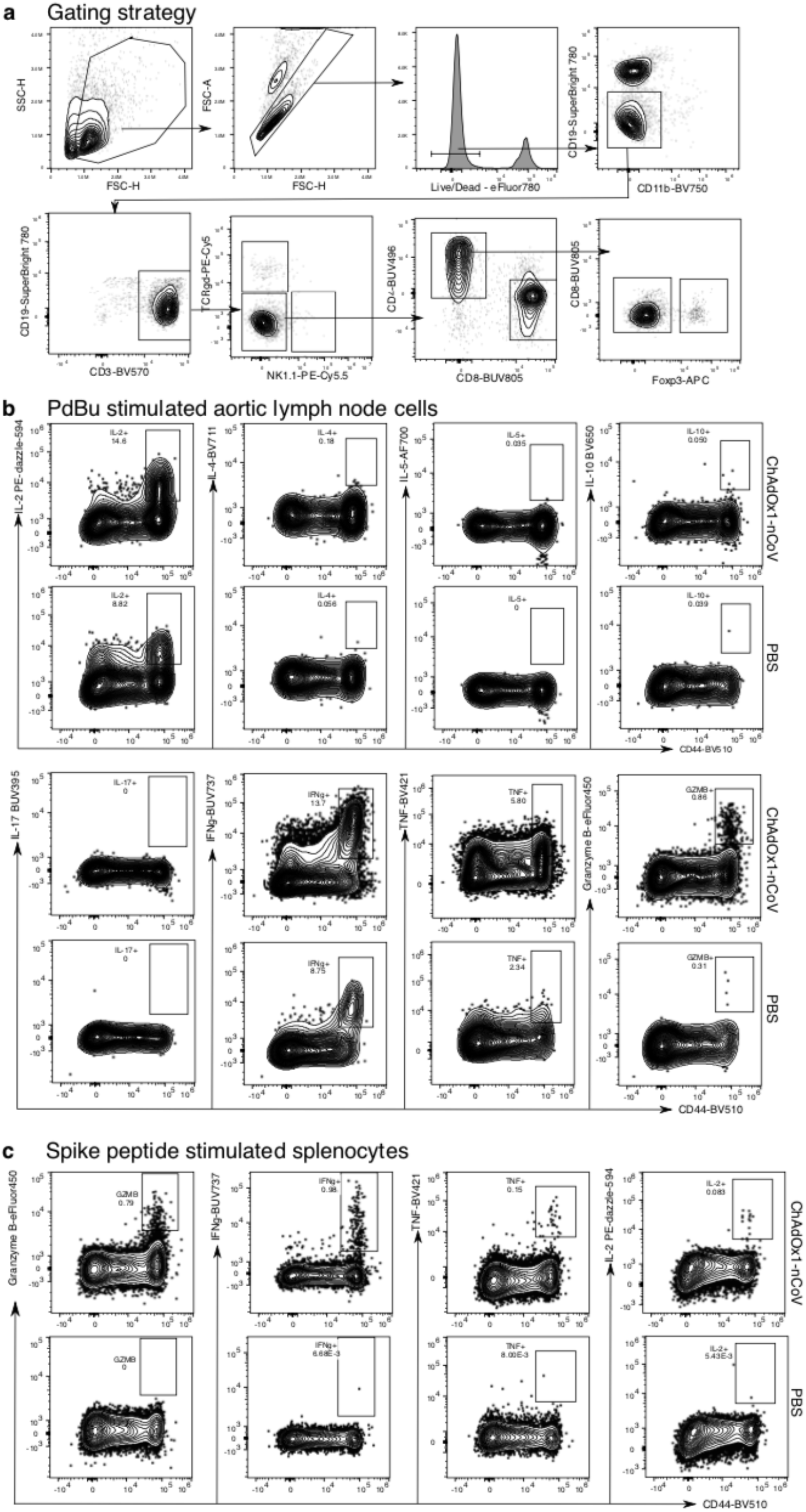
Gating strategy CD8 T cell cytokine and granzyme B staining. Flow cytometric contour plots showing the gating strategy to identify T cell subsets (**a**), cytokine and granzyme B production in CD8^+^ aortic lymph node T cells six hours after PdBu/Ionomycin restimulation (**b**), and in SARS-CoV-2 peptide restimulated CD8^+^ splenocytes (**c**).

**Supplementary Figure 5.**
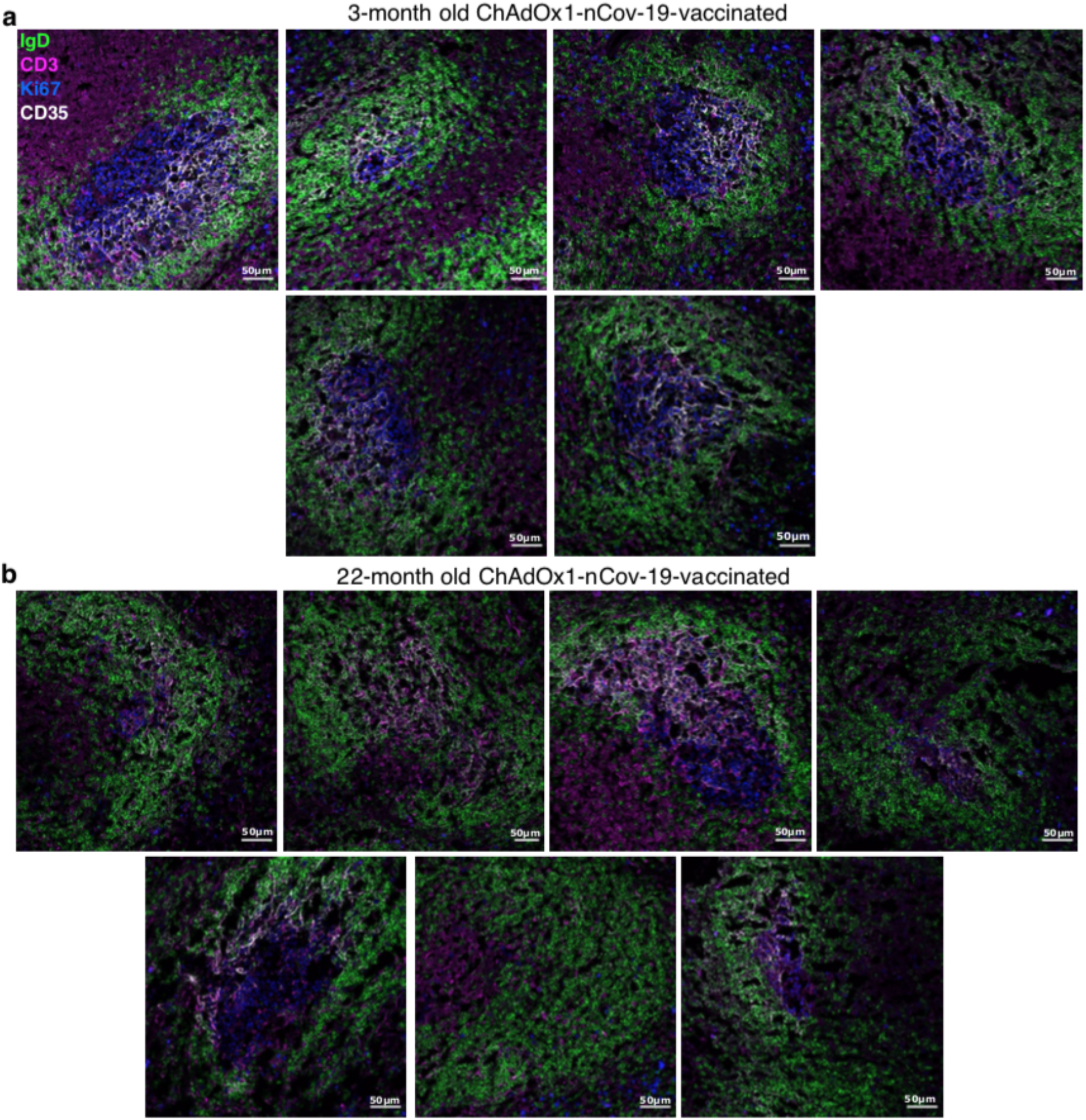
Germinal centre images nine days after ChAdOx1 nCoV-19 immunisation in adult and aged mice. Confocal images of the spleen of ChAdOx1 nCoV-19 immunised 3-month-old (**a**) and 22-month-old (**b**) mice. 10 μm spleen sections were stained with anti-IgD (green), anti-CD3 (pink), anti-Ki67 (blue) and anti-CD35 (white) antibodies. Each image is from a different mouse. The scale bars indicate 50μm.

## References

1. N. Zhu et al., A Novel Coronavirus from Patients with Pneumonia in China, 2019. N Engl J Med 382, 727–733 (2020).

2. F. Wu et al., A new coronavirus associated with human respiratory disease in China. Nature 579, 265–269 (2020).

3. M. Hoffmann et al., SARS-CoV-2 Cell Entry Depends on ACE2 and TMPRSS2 and Is Blocked by a Clinically Proven Protease Inhibitor. Cell 181, 271–280 e278 (2020).

4. Z. Wu, J. M. McGoogan, Characteristics of and Important Lessons From the Coronavirus Disease 2019 (COVID-19) Outbreak in China: Summary of a Report of 72314 Cases From the Chinese Center for Disease Control and Prevention. JAMA, (2020).

5. C. Wu et al., Risk Factors Associated With Acute Respiratory Distress Syndrome and Death in Patients With Coronavirus Disease 2019 Pneumonia in Wuhan, China. JAMA Intern Med, (2020).

6. G. Grasselli et al., Baseline Characteristics and Outcomes of 1591 Patients Infected With SARS-CoV-2 Admitted to ICUs of the Lombardy Region, Italy. JAMA, (2020).

7. B. Greenwood, The contribution of vaccination to global health: past, present and future. Philos Trans R Soc Lond B Biol Sci 369, 20130433 (2014).

8. M. Doherty, P. Buchy, B. Standaert, C. Giaquinto, D. Prado-Cohrs, Vaccine impact: Benefits for human health. Vaccine 34, 6707–6714 (2016).

9. N. Lurie, M. Saville, R. Hatchett, J. Halton, Developing Covid-19 Vaccines at Pandemic Speed. N Engl J Med 382, 1969–1973 (2020).

10. H. R. Sharpe et al., The early landscape of coronavirus disease 2019 vaccine development in the UK and rest of the world. Immunology 160, 223–232 (2020).

11. T. M. Govaert et al., The efficacy of influenza vaccination in elderly individuals. A randomized double-blind placebo-controlled trial. Jama 272, 1661–1665 (1994).

12. A. Ortqvist et al., Randomised trial of 23-valent pneumococcal capsular polysaccharide vaccine in prevention of pneumonia in middle-aged and elderly people. Swedish Pneumococcal Vaccination Study Group. Lancet 351, 399–403 (1998).

13. K. Stiasny, J. H. Aberle, M. Keller, B. Grubeck-Loebenstein, F. X. Heinz, Age affects quantity but not quality of antibody responses after vaccination with an inactivated flavivirus vaccine against tick-borne encephalitis. PLoS One 7, e34145 (2012).

14. E. A. Burns, L. G. Lum, G. L’Hommedieu, J. S. Goodwin, Specific humoral immunity in the elderly: in vivo and in vitro response to vaccination. J Gerontol 48, B231–236 (1993).

15. C. A. DiazGranados et al., Efficacy of high-dose versus standard-dose influenza vaccine in older adults. N Engl J Med 371, 635–645 (2014).

16. S. Squarcione, S. Sgricia, L. R. Biasio, E. Perinetti, Comparison of the reactogenicity and immunogenicity of a split and a subunit-adjuvanted influenza vaccine in elderly subjects. Vaccine 21, 1268–1274 (2003).

17. V. Vanhooren, C. Libert, The mouse as a model organism in aging research: usefulness, pitfalls and possibilities. Ageing Res Rev 12, 8–21 (2013).

18. J. J. Goronzy, C. M. Weyand, Mechanisms underlying T cell ageing. Nat Rev Immunol 19, 573–583 (2019).

19. D. Frasca, B. B. Blomberg, D. Garcia, S. R. Keilich, L. Haynes, Age-related factors that affect B cell responses to vaccination in mice and humans. Immunol Rev 296, 142–154 (2020).

20. C. E. Gustafson, C. Kim, C. M. Weyand, J. J. Goronzy, Influence of immune aging on vaccine responses. J Allergy Clin Immunol 145, 1309–1321 (2020).

21. M. Stebegg et al., Rejuvenating conventional dendritic cells and T follicular helper cell formation after vaccination. Elife 9, (2020).

22. H. I. Nakaya et al., Systems Analysis of Immunity to Influenza Vaccination across Multiple Years and in Diverse Populations Reveals Shared Molecular Signatures. Immunity 43, 1186–1198 (2015).

23. X. Yang, J. Stedra, J. Cerny, Relative contribution of T and B cells to hypermutation and selection of the antibody repertoire in germinal centers of aged mice. J Exp Med 183, 959–970 (1996).

24. J. S. Lefebvre et al., The aged microenvironment contributes to the age-related functional defects of CD4 T cells in mice. Aging Cell 11, 732–740 (2012).

25. T. M. Govaert et al., The efficacy of influenza vaccination in elderly individuals. A randomized double-blind placebo-controlled trial. JAMA 272, 1661–1665 (1994).

26. R. Praditsuwan, P. Assantachai, C. Wasi, P. Puthavatana, U. Kositanont, The efficacy and effectiveness of influenza vaccination among Thai elderly persons living in the community. J Med Assoc Thai 88, 256–264 (2005).

27. S. L. Baldwin et al., Improved Immune Responses in Young and Aged Mice with Adjuvanted Vaccines against H1N1 Influenza Infection. Front Immunol 9, 295 (2018).

28. V. Demicheli, T. Jefferson, E. Ferroni, A. Rivetti, C. Di Pietrantonj, Vaccines for preventing influenza in healthy adults. Cochrane Database Syst Rev 2, CD001269 (2018).

29. I. F. Hung et al., Immunogenicity of intradermal trivalent influenza vaccine with topical imiquimod: a double blind randomized controlled trial. Clin Infect Dis 59, 1246–1255 (2014).

30. D. L. Hill et al., The adjuvant GLA-SE promotes human Tfh cell expansion and emergence of public TCRbeta clonotypes. J Exp Med 216, 1857–1873 (2019).

31. N. van Doremalen et al., ChAdOx1 nCoV-19 vaccine prevents SARS-CoV-2 pneumonia in rhesus macaques. Nature, (2020).

32. S. P. Graham et al., Evaluation of the immunogenicity of prime-boost vaccination with the replication-deficient viral vectored COVID-19 vaccine candidate ChAdOx1 nCoV-19. npj Vaccines 5, (2020).

33. P. M. Folegatti et al., Safety and immunogenicity of the ChAdOx1 nCoV-19 vaccine against SARS-CoV-2: a preliminary report of a phase 1/2, single-blind, randomised controlled trial. Lancet, (2020).

34. V. Manolova et al., Nanoparticles target distinct dendritic cell populations according to their size. Eur J Immunol 38, 1404–1413 (2008).

35. M. Guilliams et al., Unsupervised High-Dimensional Analysis Aligns Dendritic Cells across Tissues and Species. Immunity 45, 669–684 (2016).

36. J. M. den Haan, S. M. Lehar, M. J. Bevan, CD8(+) but not CD8(-) dendritic cells cross-prime cytotoxic T cells in vivo. J Exp Med 192, 1685–1696 (2000).

37. A. Mildner, S. Jung, Development and function of dendritic cell subsets. Immunity 40, 642–656 (2014).

38. I. C. MacLennan et al., Extrafollicular antibody responses. Immunol Rev 194, 8–18 (2003).

39. C. G. Vinuesa, M. A. Linterman, D. Yu, I. C. MacLennan, Follicular Helper T Cells. Annu Rev Immunol 34, 335–368 (2016).

40. D. Frasca, E. Van der Put, R. L. Riley, B. B. Blomberg, Reduced Ig class switch in aged mice correlates with decreased E47 and activation-induced cytidine deaminase. J Immunol 172, 2155–2162 (2004).

41. D. Frasca et al., Aging down-regulates the transcription factor E2A, activation-induced cytidine deaminase, and Ig class switch in human B cells. J Immunol 180, 5283–5290 (2008).

42. D. Frasca et al., Intrinsic defects in B cell response to seasonal influenza vaccination in elderly humans. Vaccine 28, 8077–8084 (2010).

43. M. J. Shlomchik, W. Luo, F. Weisel, Linking signaling and selection in the germinal center. Immunol Rev 288, 49–63 (2019).

44. L. Mesin, J. Ersching, G. D. Victora, Germinal Center B Cell Dynamics. Immunity 45, 471–482 (2016).

45. F. J. Weisel, G. V. Zuccarino-Catania, M. Chikina, M. J. Shlomchik, A Temporal Switch in the Germinal Center Determines Differential Output of Memory B and Plasma Cells. Immunity 44, 116–130 (2016).

46. B. Zheng, S. Han, Y. Takahashi, G. Kelsoe, Immunosenescence and germinal center reaction. Immunol Rev 160, 63–77 (1997).

47. J. S. Tsang et al., Global analyses of human immune variation reveal baseline predictors of postvaccination responses. Cell 157, 499–513 (2014).

48. F. C. Zhu et al., Immunogenicity and safety of a recombinant adenovirus type-5-vectored COVID-19 vaccine in healthy adults aged 18 years or older: a randomised, double-blind, placebo-controlled, phase 2 trial. Lancet, (2020).

49. P. J. M. Brouwer et al., Potent neutralizing antibodies from COVID-19 patients define multiple targets of vulnerability. Science, (2020).

50. C. Kreer et al., Longitudinal Isolation of Potent Near-Germline SARS-CoV-2-Neutralizing Antibodies from COVID-19 Patients. Cell, (2020).

51. E. Seydoux et al., Analysis of a SARS-CoV-2-Infected Individual Reveals Development of Potent Neutralizing Antibodies with Limited Somatic Mutation. Immunity 53, 98–105 e105 (2020).

52. P. T. Sage, C. L. Tan, G. J. Freeman, M. Haigis, A. H. Sharpe, Defective TFH Cell Function and Increased TFR Cells Contribute to Defective Antibody Production in Aging. Cell Rep 12, 163–171 (2015).

53. D. Botta et al., Dynamic regulation of T follicular regulatory cell responses by interleukin 2 during influenza infection. Nat Immunol 18, 1249–1260 (2017).

54. A. E. Denton et al., Intrinsic defects in lymph node stromal cells underpin poor germinal center responses during aging. bioRxiv, (2020).

55. M. R. Bassi et al., CD8+ T cells complement antibodies in protecting against yellow fever virus. J Immunol 194, 1141–1153 (2015).

56. J. D. Brien, J. L. Uhrlaub, A. Hirsch, C. A. Wiley, J. Nikolich-Zugich, Key role of T cell defects in age-related vulnerability to West Nile virus. J Exp Med 206, 2735–2745 (2009).

57. C. C. Clay et al., Severe acute respiratory syndrome-coronavirus infection in aged nonhuman primates is associated with modulated pulmonary and systemic immune responses. Immun Ageing 11, 4 (2014).

58. J. E. McElhaney et al., T cell responses are better correlates of vaccine protection in the elderly. J Immunol 176, 6333–6339 (2006).

59. D. C. Powers, R. B. Belshe, Effect of age on cytotoxic T lymphocyte memory as well as serum and local antibody responses elicited by inactivated influenza virus vaccine. J Infect Dis 167, 584–592 (1993).

60. J. E. McElhaney et al., Granzyme B: a marker of risk for influenza in institutionalized older adults. Vaccine 19, 3744–3751 (2001).

61. M. D. Dicks et al., A novel chimpanzee adenovirus vector with low human seroprevalence: improved systems for vector derivation and comparative immunogenicity. PLoS One 7, e40385 (2012).

62. M. Mahler Convenor et al., FELASA recommendations for the health monitoring of mouse, rat, hamster, guinea pig and rabbit colonies in breeding and experimental units. Lab Anim 48, 178–192 (2014).

63. E. Pasciuto et al., Microglia Require CD4 T Cells to Complete the Fetal-to-Adult Transition. Cell 182, 625–640 e624 (2020).

